# Structural basis for binding of Smaug to the GPCR Smoothened and to the germline inducer Oskar

**DOI:** 10.1101/2023.02.19.529116

**Authors:** Jana Kubíková, Gabrielė Ubartaitė, Jutta Metz, Mandy Jeske

## Abstract

*Drosophila* Smaug and its orthologs comprise a family of mRNA repressor proteins that exhibit various functions during animal development. Smaug proteins contain a characteristic RNA-binding sterile-α motif (SAM) domain and a conserved but uncharacterized N-terminal domain (NTD). Here, we resolved the crystal structure of the NTD of the human SAM domain-containing protein 4A (SAMD4A, a.k.a. Smaug1) to 2.0 Å resolution, which revealed its composition of a homodimerization D-subdomain and a subdomain with similarity to a PHAT domain. Furthermore, we show that *Drosophila* Smaug directly interacts with the *Drosophila* germline inducer Oskar and with the Hedgehog signaling transducer Smoothened through its D-PHAT domain. We determined the crystal structure of the D-PHAT domain of Smaug in complex with a Smoothened α-helical peptide to 1.61 Å resolution. The peptide binds within a groove that is formed by both the D- and PHAT subdomains. Structural modeling supported by experimental data suggested that an α-helix within the disordered region of Oskar binds to the D-PHAT domain in a mode similar to Smoothened. Together, our data uncover the N-terminal D-PHAT domain of Smaug as peptide-binding domain.

## INTRODUCTION

During oogenesis and early embryogenesis, maternally deposited mRNAs and proteins determine the developmental program in many animals. Over this period of time, the genome is transcriptionally silent, and the expression of genes is regulated at the post-transcriptional level by the modulation of mRNA translation, localization, and stability (Lasko, 2011; Lazzaretti & Bono, 2017; Schoenberg & Maquat, 2012). Regulated mRNAs often carry specific *cis*-elements which are recognized by *trans*-acting RNA-binding proteins or microRNAs (miRNAs) that regulate a transcript’s fate (Fabian & Sonenberg, 2012; Lee *et al*, 2020).

*Drosophila* Smaug and its metazoan orthologs comprise a family of mRNA repressor proteins that contain a characteristic sterile-α motif (SAM) domain, two ‘Smaug similarity regions’ SSR1 and SSR2 within their N-terminal segment, and a pseudo-HEAT-repeat analogous topology (PHAT) domain C-terminally to the SAM domain **(Figure 1A**). The SAM domain is one of the most abundant protein-protein interaction domains (Thanos *et al*, 1999; Schultz *et al*, 1997; Qiao & Bowie, 2005). In Smaug orthologs, however, it serves as RNA binding domain and recognizes mRNA targets through binding to defined stem-loop structures, designated ‘Smaug recognition elements’ (SREs). (Amadei *et al*, 2015; Aviv *et al*, 2003; Baez & Boccaccio, 2005; Green *et al*, 2003; Niu *et al*, 2017; Johnson & Donaldson, 2006; Aviv *et al*, 2006; Oberstrass *et al*, 2006). The SSR2 appears limited to metazoan Smaug orthologs, while the SSR1 is present also in various other proteins, including F-box proteins, and was shown to form homodimers (Tang *et al*, 2007). The functions of the SSR2 and the PHAT domain remained unclear.

**Figure 1.**
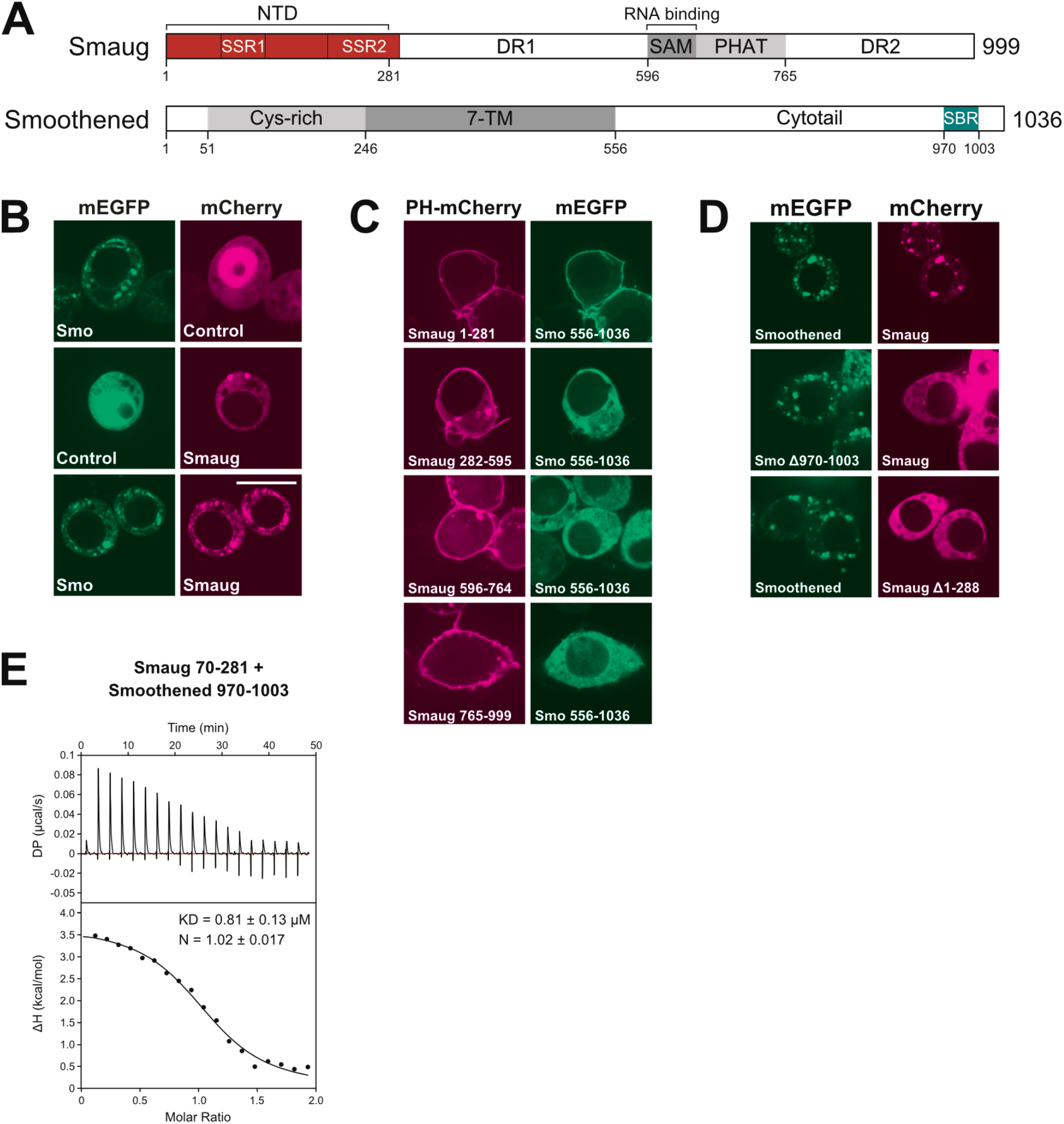
The Smaug - Smoothened interaction. (A) Domain organization of *Drosophila* Smaug and Smoothened proteins. SSR, Smaug similarity region; NTD, N-terminal domain; DR, disordered region; TM, transmembrane; SBR, Smaug-binding region. (B) ReLo assays showing a colocalization between Smaug and Smoothened. Controls are plasmids without an insert. The scale bar is 10 μm. (C) Of the four Smaug fragments indicated, only the NTD (Smaug 1-281) interacts with the Smoothened cytotail (Smo 556-1036) in the ReLo assay. The scale bar is 10 μm. (D) The Smoothened deletion and the Smaug deletion are unable to interact with Smaug and Smoothened, respectively, in the ReLo assay. The scale bar is 10 μm. (E) Isothermal titration calorimetry using Smaug NTD (Smaug 70-281) and a Smoothened peptide (Smo 970-1003).

In animals, members of the Smaug protein family exhibit various functions during development. Mammals express two Smaug-related paralogs named SAM-domain-containing protein 4A (SAMD4A, a.k.a. Smaug1) and SAMD4B (a.k.a. Smaug2). A SAMD4A mis-sense mutation in mice causes a lean phenotype and these animals develop kyphosis associated with myopathy and adipocyte defects, and show delayed bone development and decreased osteogenesis (Chen *et al*, 2014b; Niu *et al*, 2017). Mouse SAMD4B, but not SAMD4A, is present in neuronal precursors of mouse embryos and inhibits neurogenesis in the embryonic cortex (Amadei *et al*, 2015).

*Drosophila* Smaug mediates translation inhibition and degradation of bulk *nanos* mRNA in the early *Drosophila* embryo, a critical step during embryonic patterning (Dahanukar & Wharton, 1996; Dahanukar *et al*, 1999; Smibert *et al*, 1996). Smaug mediates its effects by recruiting the eIF4E-binding translation inhibitor Cup and the CCR4-NOT deadenylase complex (Götze *et al*, 2017; Jeske *et al*, 2011; Nelson *et al*, 2004; Semotok *et al*, 2005; Zaessinger *et al*, 2006; Pekovic *et al*, 2022). Several other RNA-binding proteins have been implicated in Smaug-mediated repression of *nanos* mRNA, including Argonaute 1 (Ago1), Trailer Hitch (Tral), and the DEAD-box RNA helicases ‘Maternal expression at 31B’ (Me31B; ortholog of human DDX6) and Belle (ortholog of human DDX3X/DDX3Y) (Götze *et al*, 2017; Pinder & Smibert, 2013).

At the posterior pole of the *Drosophila* embryo, a small fraction of localized *nanos* mRNA escapes Smaug-mediated repression through the activity of Oskar protein (Dahanukar & Wharton, 1996; Dahanukar *et al*, 1999; Jeske *et al*, 2011; Zaessinger *et al*, 2006). Restriction of Nanos synthesis to the posterior pole gives rise to a Nanos protein gradient that determines the position of abdominal structures in the early embryo (Wang & Lehmann, 1991; Wang *et al*, 1994; Gavis & Lehmann, 1992, 1994). Genetic data revealed that Smaug overexpression antagonizes the Oskar-dependent activation of *nanos* mRNA in embryos (Dahanukar *et al*, 1999). Translation derepression of *nanos* mRNA can be recapitulated *in vitro* by addition of recombinant Oskar protein to a cell-free system that is active in both the translation repression and deadenylation of a *nanos* 3’ UTR-containing reporter mRNA (Jeske *et al*, 2011). Furthermore, pull-down experiments using lysates prepared from early *Drosophila* embryos demonstrated that in the presence of Oskar, Smaug is unable to bind to *nanos* mRNA (Jeske *et al*, 2011; Zaessinger *et al*, 2006). Yet, the molecular mechanisms underlying the Oskar-dependent derepression of *nanos* mRNA at the posterior pole of the embryo remain unclear.

In addition to *nanos* mRNA, Smaug acts on many other target transcripts during early embryogenesis and plays a key role in the maternal-to-zygotic transition (MZT) of gene expression, syncytial cell cycle control, blastoderm cellularization, and gastrulation (Benoit *et al*, 2009; Chen *et al*, 2014a; Siddiqui *et al*, 2012; Tadros *et al*, 2007). Smaug protein is highly abundant during the first three hours of embryogenesis (Cao *et al*, 2020; Dahanukar *et al*, 1999; Smibert *et al*, 1999). Later in embryogenesis, with the onset of zygotic transcription, Smaug protein is targeted by a Skp/Cullin/F-box-containing (SCF) ubiquitin E3 ligase complex and is subsequently degraded by the ubiquitin-proteasome system, a process required for an orderly MZT (Cao *et al*, 2022, 2020).

Through studies in cultured *Drosophila* CI8 cells, in *Drosophila* wing imaginal discs, and in *Drosophila* wings, Smaug has been linked to the Hedgehog (HH) signaling pathway (Bruzzone *et al*, 2020). It was shown that Smaug binds to Smoothened, a protein that is structurally similar to Frizzled-type G-protein-coupled receptors (GPCRs) and essential for the transduction of the HH signal (Ingham & McMahon, 2001). The recruitment of Smaug to Smoothened was proposed to result in phosphorylation of Smaug by the protein kinase Fused (Bruzzone *et al*, 2020). Whether the Smaug - Smoothened interaction and the phosphorylation of Smaug play a role during early embryonic development has not been addressed.

Here, we performed a protein-protein interaction screen and confirmed that Oskar and Smoothened directly bind Smaug. Both proteins associated with the previously uncharacterized N-terminal domain (NTD) of Smaug comprising both SSR1 and SSR2. We have solved the crystal structures of the NTD of the human Smaug ortholog SAMD4A alone and of the NTD of *Drosophila* Smaug in complex with a Smoothened peptide. The crystal structures revealed that the NTD is composed of a dimerization (D) subdomain and a PHAT subdomain, which interact and jointly form a groove that is bound by the Smoothened peptide. Furthermore, we identified a predicted α-helix within the disordered region of Oskar that binds to the NTD of Smaug, and structural modeling supported by experimental data suggested that the complex is structurally similar to the Smaug-Smoothened complex. Together, our data uncover the structural basis for the complex formation between an RNA-binding protein and a signaling protein, and suggest a conserved function of the NTD of Smaug proteins as peptide-binding domain.

## RESULTS

### Smaug directly binds to Smoothened

To analyze protein interactions between proteins, we have recently developed ReLo, a cell culture-based protein-protein interaction assay that is based on a subcellular translocation readout (Salgania *et al*, 2022). Our previous analyses strongly suggested that interactions identified with the ReLo assay are direct (Salgania *et al*, 2022). Furthermore, we have shown that the ReLo assay is particularly suitable for the identification and characterization of direct interactions to proteins that are large and poorly accessible for biochemical and biophysical studies, such as Smaug. In the ReLo assay, the two proteins to be tested for an interaction are fused to red (mCherry) or green (EGFP) fluorescent proteins, coexpressed in *Drosophila* Schneider 2R+ (S2R+) cells and their subcellular localization is analyzed by confocal fluorescence microscopy. Importantly, the bait protein carries a membrane anchoring domain, and its interaction with a prey protein is visualized by the subcellular relocalization of the prey protein towards the membrane to which the bait protein is anchored.

To assess protein interactions with Smaug we used an EGFP-Smaug construct that carried an N-terminal fusion to a PH domain (Salgania *et al*, 2022), which directed the localization of Smaug to the plasma membrane. With the ReLo assay, we have previously evaluated the interaction between Smaug and the six core subunits of the CCR4-NOT deadenylase complex, which is responsible for the deadenylation of *nanos* mRNA, and have identified the NOT3 subunit as Smaug-binding protein (Pekovic *et al*, 2022). Here, we tested interactions between Smaug and additional proteins involved in *nanos* mRNA repression, including Ago1 and Ago2 (Pinder and Smibert, 2013), Aubergine and Ago3 (Barckmann *et al*, 2015; Rouget *et al*, 2010), Cup (Nelson *et al*, 2004) as well as Me31B, Tral, and Belle (Götze *et al*, 2017; Jeske *et al*, 2011). However, we did not observe an interaction between Smaug and any of these factors **(Supplemental Figure 1A).** In addition to the pairwise testing, the Smaug-Cup interaction was also tested in the presence of eIF4E and/or the translation control element (TCE) of *nanos* mRNA, which contains the Smaug recognition elements (SRE). Again, no interaction between Smaug and Cup was observed (**Supplemental Figure 1B**).

Recent findings indicated a direct interaction between Smaug and Smoothened (Bruzzone *et al*, 2020). Smoothened is a membrane protein that localizes to distinct cytoplasmic vesicles and as such we were able to confirm the Smoothened-Smaug interaction in the ReLo assay by asking if Smoothened localizes Smaug to these vesicles (**Figures 1A and 1B**). We next set out to map the regions of Smaug and Smoothened that are involved in the interaction. Smoothened consists of an N-terminal extracellular cysteine-rich domain (CRD; aa 51-246), a central seven-transmembrane domain (7-TM; aa 247-555), and a C-terminal ‘cytotail’ (aa 556-1036), which is a long cytoplasmic and predominantly disordered region **(Figure 1A).** As the cytotail has been previously reported to bind Smaug (Bruzzone et al., 2020), this fragment of Smoothened was tested in combination with the N-terminal part of Smaug (NTD; aa 1-281), the disordered region (DR1, aa 282-595), the SAM-PHAT domain combination (aa 596-765), or the C-terminal disordered region (DR2, aa 766-999). The results revealed that the Smoothened cytotail bound to the NTD of Smaug **(Figure 1C),** which is consistent with previous data (Bruzzone *et al*, 2020). Using ReLo, we further confirmed previous data that Smaug bound to a small region within the cytotail of Smoothened spanning amino acid residues 958-1003 (Bruzzone *et al*, 2020), and refined the mapping to a predicted α–helix covering residues 970-1003 of Smoothened (**Figure 1D**). By isothermal titration calorimetry (ITC), the K_D_ of the complex formed by Smaug 70-281 and a synthetic Smo 970-1003 peptide was determined as 0.8 μM, and the complex was found to have a 1:1 stoichiometry **(Figure 1E**). In following experiments, we refer to Smoothened 970-1003 as the Smaug binding region (SBR).

### Crystal structure of the D-PHAT domain of human SAMD4A

We sought to determine the three-dimensional structure of the Smoothened-binding NTD of *Drosophila* Smaug by X-ray crystallography. Unfortunately, none of the soluble Smaug-NTD variants tested containing or lacking N-terminal truncations (Δ1-36, Δ1-69) or regions predicted to be disordered (Δ156-196, Δ175-184, Δ159-184) crystallized. As an alternative, we purified and crystallized the NTD (aa 2-156) of the human Smaug ortholog SAMD4A **(Figure 2A)** as a hexahistidine (His)-tag fusion. The crystals obtained diffracted to 1.62 Å resolution, and additional diffraction experiments using selenomethionine-derivative crystals allowed us to solve the structure by native phasing methods **(Supplemental Table 1).**

**Figure 2.**
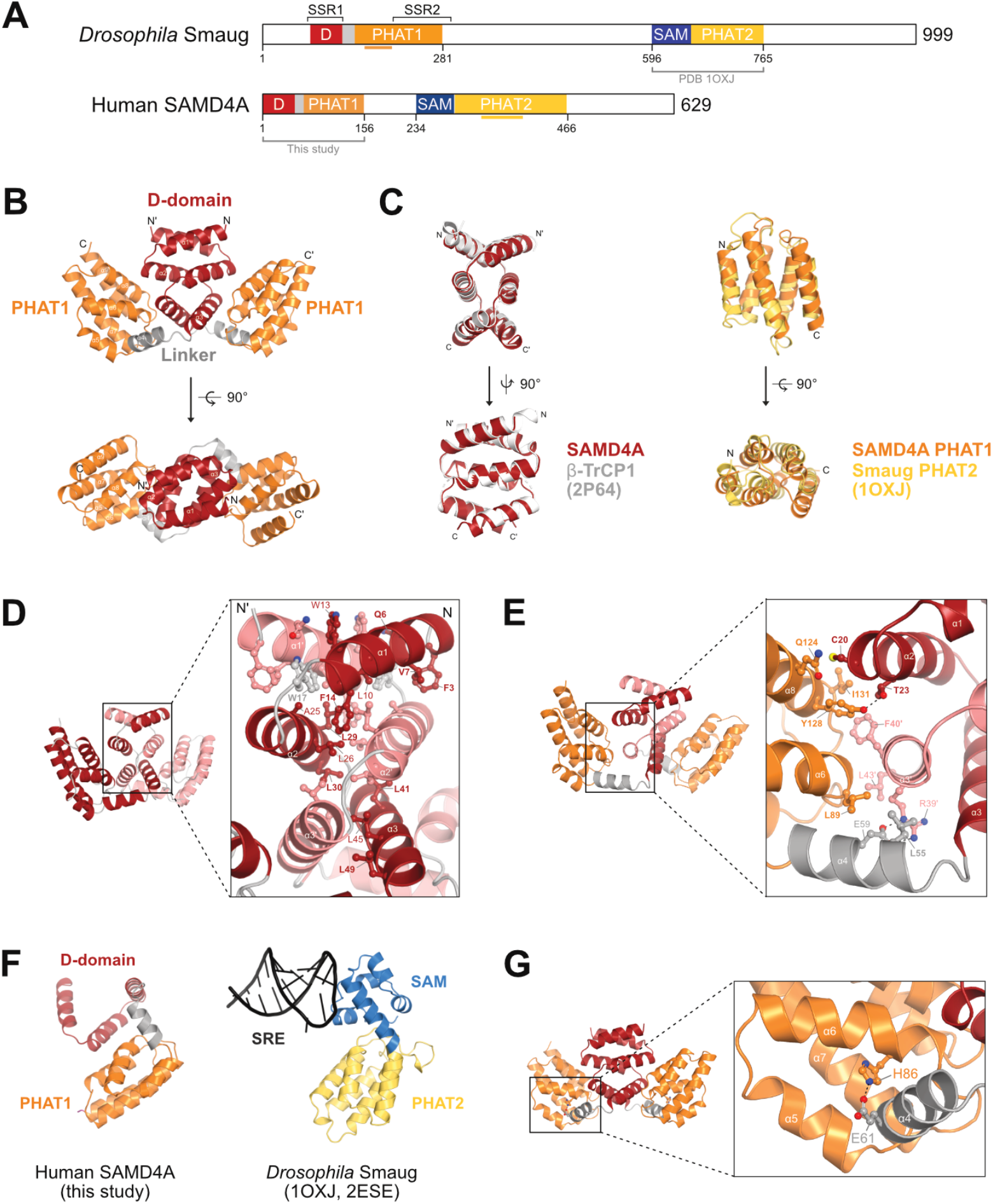
Crystal structure of the NTD of human SAMD4A. (A) Domain organization of *Drosophila* Smaug and human SAMD4A proteins. The horizontal lines below the protein schemes indicate insertions into the respective PHAT domains. (B) Crystal structure of the NTD dimer of the human SAMD4A. The D-subdomain, the PHAT1-subdomain and the linker α-helix are indicated. (C) Structural superimposition of the D-subdomains of SAMD4A onto the D-domain of β-TrCP (left panels) and of the PHAT1 domain of SAMD4A onto the PHAT2 domain of Smaug (right panels). (D) Details of the D-subdomain dimer interface of SAMD4A. Amino acid residues are labeled for one protein chain only. Residues in bold are 100% conserved across animals. (E) Details of the interface formed by the D- and the PHAT1 subdomains. Residues in bold are 100% conserved across animals. (F) Structural comparison between the D-PHAT1 domain of SAMD4A and the SAM-PHAT2 domain of Smaug. SRE, Smaug recognition elements. The model of the SAM-PHAT2 domain - SRE complex was prepared by superimposition of the two crystals structures indicated. (G) Structural detail indicating the position of the H86 residue, which is mutated in the *spmd* mutation.

The crystal structure revealed that the SAMD4A-NTD forms a homodimer, and each monomer is composed of two subdomains **(Figure 2B).** The N-terminal subdomain is composed of three α–helices that exhibit a zig-zag arrangement and has previously been predicted to form a dimerization domain similar to the D-domain of the β-Transducin repeat-containing protein 1 (β-TrCP1) (Tang *et al*, 2007). In fact, a search with the protein structure comparison server DALI (Holm & Rosenström, 2010) revealed that the D-subdomain of SAMD4A is structurally most similar to the D-domain of β-TrCP1. Both D-domain dimers align with a root mean square deviation (RMSD) of 0.928 Å over 328 aligned atoms **(Figure 2C, left panel).** Within the dimer, the two D-subdomains are parallelly oriented and extensively interlaced to establish a superhelical tertiary structure. The hydrophobic dimer interface is highly conserved and buries a surface area of 2645 Å^2^ as determined using the PISA server (Krissinel & Henrick, 2007) **(Figure 2D and Supplemental Figure 2).** Attempts to interfere with dimerization through point mutagenesis led to insoluble protein in recombinant expression experiments (**data not shown**).

The C-terminal subdomain of the SAMD4A-NTD is composed of a bundle of five α–helices (α5-9), which is connected with the D-subdomain through a linker α–helix (α4) **(Figure 2B).** DALI search revealed that the five-helical bundle of SAMD4A is structurally most similar to the previously described PHAT domain in the middle part of *Drosophila* Smaug (Green *et al*, 2003) **(Figure 2A).** Thus, the crystal structure of the SAMD4-NTD led to discovery that both Smaug and SAMD4A contain not one but two PHAT domains: PHAT1 is connected to a D-subdomain and located in the NTD, whereas PHAT2 is connected to an RNA-binding SAM domain and located in the middle part of the protein. Sequence alignment and secondary structure prediction analysis revealed that, depending on the protein, PHAT domains are either compact or carry unstructured insertions: The PHAT1 domain of SAMD4A is compact, while the PHAT1 domain of Smaug contains two unstructured insertions; in contrast, the PHAT2 domain of Smaug is compact, while the PHAT2 domain of SAMD4A carries an unstructured insertion **(Supplemental Figures 3A and 3B).** The crystal structures of SAMD4A PHAT 1 and Smaug PHAT2 aligned with an RMSD of 2.87 Å over 88 residues **(Figure 2C).** The *Saccharomyces cerevisiae* ortholog Vts1 lacks both PHAT domains **(Supplemental Figure 3C).**

In the SAMD4A-NTD structure, the PHAT1-domain contacts both monomers of the D-subdomain dimer - the one in the same polypeptide chain and its dimerizing partner molecule **(Figure 2E).** Furthermore, the residues of interface between the D and PHAT1 subdomains are highly conserved (**Supplemental Figure 2**). This suggested that the PHAT1 subdomain folds back to the D-subdomain only after the dimer has been established. A rigid arrangement has also been discussed previously for the relative position of the PHAT2 domain to the SAM domain of Smaug (Green *et al*, 2003) **(Figure 2F).**

A point mutation in mouse SAMD4A causes a *supermodel* (*spmd*) phenotype with mice being resistant to obesity induced by a high fat diet, and displaying leanness and myopathy (Chen *et al*, 2014b). The *spmd* mutation (H86P) maps to the NTD of SAMD4A, and we analyzed the mutation using our structure of the human SAMD4A-NTD. With the exception of two residues, the sequences of the NTDs (aa 1-155) of human and mouse SAMD4A are identical **(Supplemental Figure 2A).** Our structure revealed that H86 resides within the α6-helix of the PHAT1 domain and is engaged in a hydrogen bond with E61 residing in the linker α4-helix **(Figure 2G).** Mutating H86 to proline caused complete insolubility of both human and mouse SAMD4A-NTD upon expression in *E. coli*, which is in stark contrast to the corresponding highly soluble wildtype constructs **(Supplemental Figure 4).** This indicates that the H86P mutation prevents proper folding of the domains. Proline residues can undergo a *cis-trans*-isomerization reaction and are well known for their impact on protein (mis)folding, for example by partial mis-isomerization or by their ability to induce kinks in α–helices (Englander & Mayne, 2014). Therefore, we assume that the *spmd* mutation causes folding problems of the SAMD4A-NTD *in vivo*, which probably leads to misfunctioning of the protein in mice.

### Structure of the NTD of Smaug in complex with the SBR of Smoothened

Next, we aimed to obtain structural information on the Smaug-Smoothened complex. The cytotail of Smoothened is conserved in Drosophilids but not across animals **(Supplemental Figure 5A)** and we have not detected any binding of human or *Drosophila* Smoothened to human SAMD4A **(Supplemental Figures 5B, 5C, 5D)**, which suggests that the Smoothened-Smaug interaction is specific to *Drosophila*. To crystallize the *Drosophila* Smaug-NTD in complex with the Smoothened SBR peptide, we designed a Smaug-NTD construct based on the SAMD4A-NTD structure, in which two regions of predicted disorder (Δ1-72 and Δ156-196) were deleted. The *Drosophila* Smaug-NTD harboring these two deletions was still able to bind to Smoothened (and Oskar, see below) **(Supplemental Figure 6).** Two strategies were pursued for crystallization: one, in which Smaug 73-281Δ156-196 was mixed with a Smo 970-1003 peptide and a second, in which the Smo 970-1003 sequence was fused to the N-terminus of Smaug 73-278Δ156-196 using a (GGS)_4_ linker. With both strategies, crystals of similar shape were obtained. Of these, the crystals of the single-chain construct diffracted best and to 2.0 Å resolution **(Supplemental Table 1).** The structure of the Smaug-Smoothened complex was solved by molecular replacement using the human SAMD4A-NTD structure as a search model. The resolved structure was composed of two molecules each of Smaug 73-274Δ156-196 and Smo 976-989; no electron density was observed for the (GGS)_4_ linker **(Figures 3A and 3B).**

**Figure 3.**
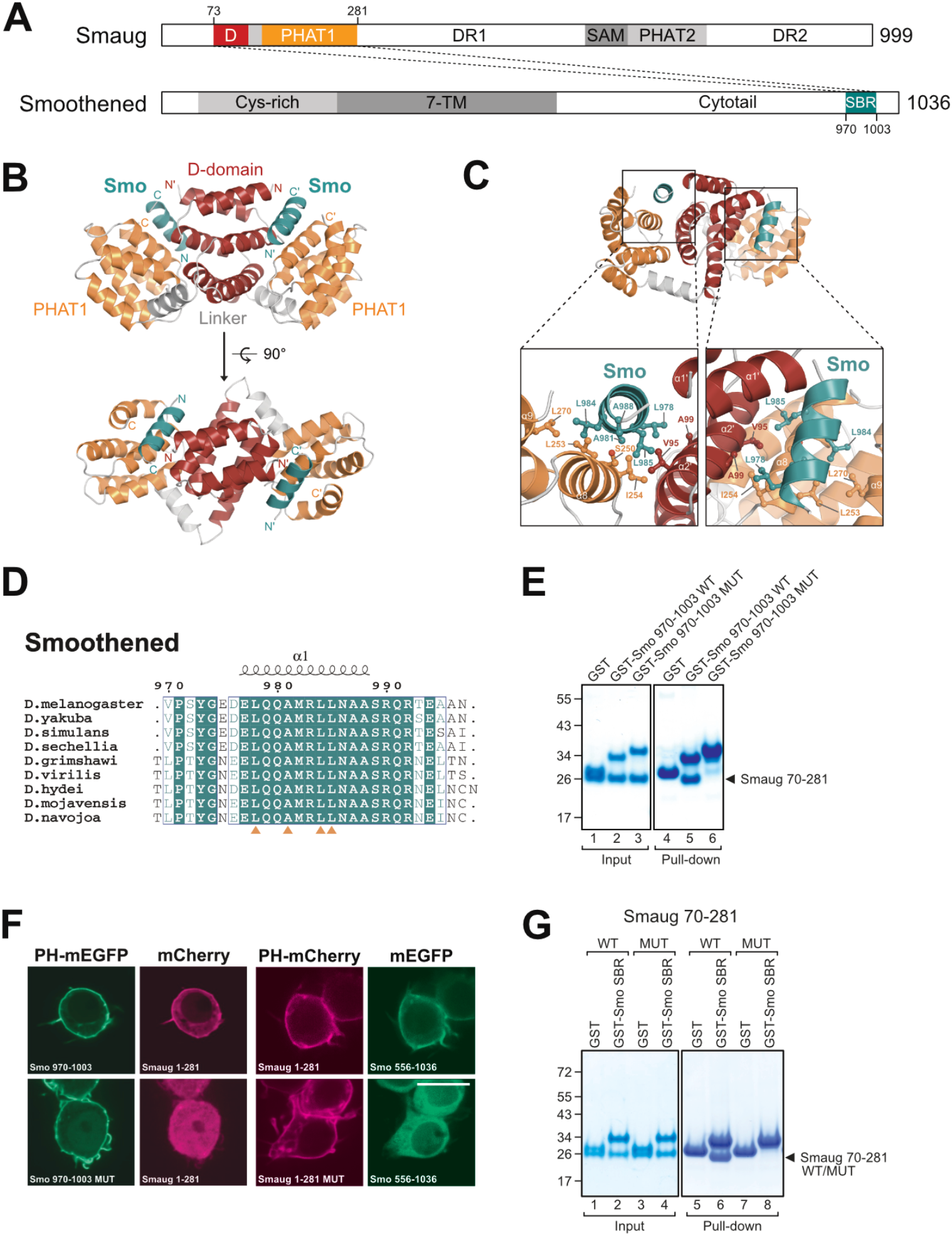
Crystals structure of Smaug-NTD in complex with Smoothened. (A) Domain organization of *Drosophila* Smaug and Smoothened proteins. NTD, N-terminal domain; DR, disordered region; TM, transmembrane; SBR, Smaug-binding region. (B) Crystal structure of the NTD of Smaug in complex with a Smoothened peptide (colored deepteal). (C) Structural detail of the Smaug - Smoothened interface. (D) Sequence alignment of the Smaug-binding region of Smoothened. Orange triangles indicate central residues involved in the Smaug interaction. (E) GST pulldown assay using GST-Smoothened 970-1003 (SBR) wildtype (WT) or L987E/L984E/L985E mutant (MUT) proteins. Protein markers in kDa are indicated on the left. (F) Mutant Smoothened SBR (MUT; L987E/L984E/L985E) did not interact with the NTD of Smaug and mutant Smaug NTD (MUT; S250E/L253E) did not interact with the Smoothened cytotail in the ReLo assay. The scale bar is 10 μm. (G) GST pulldown assay using Smaug NTD (70-281) wildtype or S250E/L253E mutant proteins and GST or GST-Smoothened SBR (970-1003). Protein markers in kDa are indicated on the left.

The Smoothened peptide forms an α–helix, which is bound to a groove created by the D-subdomain dimer and the PHAT subdomain. The interface area of the Smaug-Smoothened complex measures 1522 Å^2^ (Krissinel & Henrick, 2007) **(Figure 3C).** The Smoothened α–helix that binds to Smaug is highly conserved across Drosophilids but not in higher animals **(Figure 3D, Supplemental Figure 5A).** The Smaug-Smoothened interface was validated by mutational analyses: The triple point mutation L978E/L984E/L985E in Smoothened interfered with Smaug binding in ReLo and GST pull-down assays **(Figures 3E and 3F).** Likewise, the S250E/L253E mutation in Smaug prevented the interaction with Smoothened in ReLo and GST pull-down assays **(Figures 3F and 3G**). Thus, our mutational analysis confirms the interface of the Smaug - Smoothened complex observed in the crystal structure.

### Oskar directly binds to Smaug

Using ReLo, we also retested the previously described interaction between Smaug and Oskar (Dahanukar *et al*, 1999; Jeske *et al*, 2011; Zaessinger *et al*, 2006) (**Figure 4A**). Translation of *oskar* mRNA from two alternative start codons results in two protein isoforms, of which Short Oskar is essential for germ cell formation and posterior patterning (Markussen *et al*, 1995). We tested Smaug interaction to the two Oskar isoforms. Long Oskar is a membrane protein (Vanzo *et al*, 2007) and localized as speckles in S2R+ cells (**Figure 4B**). Thus, we did not use an additional membrane anchor to test the Smaug - Long Oskar interaction in the ReLo assay. When Long Oskar and Smaug were coexpressed, we did not observe a relocalization of Smaug **(Figure 4B).** Short Oskar localizes to the nucleus in S2R+ cells and a fusion to the PH domain does not efficiently redirect/anchor Short Oskar to the plasma membrane (Salgania *et al*, 2022). Therefore, to assess protein interactions to Short Oskar, it carried an N-terminal fusion to OST4, a small membrane protein that directed Short Oskar localization to the endoplasmic reticulum (ER). Using OST4-anchored Short Oskar, we have previously confirmed the known direct interaction with the DEAD-box RNA helicase Vasa **(Supplemental Figure 7A)** (Salgania *et al*, 2022; Jeske *et al*, 2017, 2015). Here, we show that OST4-Short Oskar interacted also with Smaug **(Figure 4B),** which is consistent with earlier observations (Dahanukar *et al*, 1999; Jeske *et al*, 2011; Zaessinger *et al*, 2006). We used additional ReLo assays to test other previously suggested binding partners of Short Oskar, including the WD40 protein Valois (a.k.a. MEP50) (Anne, 2010), the elF4E-binding protein Cup (Ottone *et al*, 2012), the actin-binding protein Lasp (Suyama *et al*, 2009), and the dsRNA-binding protein Staufen (Breitwieser *et al*, 1996). However, none of these proteins relocalized with Short Oskar **(Supplemental Figure 7B).**

**Figure 4.**
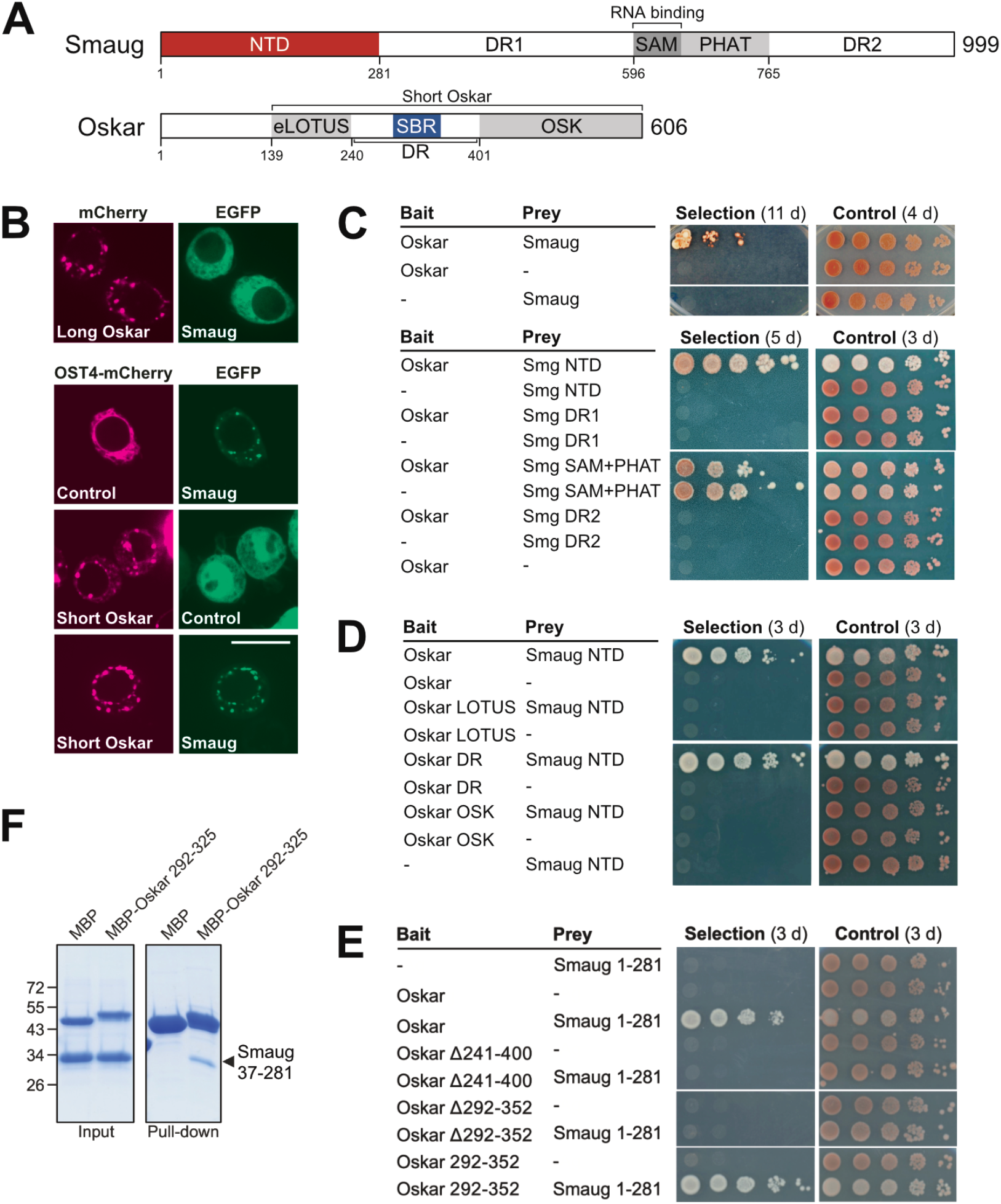
The Smaug - Short Oskar interaction. (A) Domain organization of Drosophila Smaug and Oskar. The Short Oskar isoform is generated by alternate translation from M139 onwards. NTD, N-terminal domain; DR, disordered region; SBR, Smaug-binding region. (B) Smaug interacts with Short Oskar but not Long Oskar in the ReLo assay. Plasmids lacking an insert serve as controls. The scale bar is 10 μm. (C) Oskar (only the short isoform is tested) and Smaug interact in split-ubiquitin yeast two-hybrid (Y2H) assays (upper panel). Short Oskar binding maps to the NTD of Smaug (lower panel). 10-fold dilutions are shown. Selection plates lacked adenine and histidine as compared to control plates. (D) The NTD of Smaug binds to the disordered region (DR) of Short Oskar in Y2H assays. 10-fold dilutions are shown. Selection plates lacked adenine and histidine as compared to control plates. (E) Oskar 292-352 (SBR) is necessary and sufficient for Smaug-NTD binding in Y2H assays. 10-fold dilutions are shown. Selection plates lacked adenine and histidine as compared to control plates. (F) MBP pull-down assay showing that MBP-Oskar 292-325 copurified the Smaug NTD (37-281).

The interaction between Smaug and Short Oskar was also visible in split-ubiquitin yeast-two hybrid (Y2H) assays **(Figure 4C, upper panel),** which we then used to map the respective protein regions that mediate the interaction. Interestingly, among the four Smaug fragments tested, it was the NTD that bound to Short Oskar **(Figure 4A and lower panel of Figure 4C)**, which is the same Smaug domain that also bound to Smoothened. Next, we examined which part of Short Oskar binds to Smaug. Short Oskar carries an extended LOTUS (eLOTUS) domain (aa 139-240) at the N-terminus, followed by a predominantly disordered region (DR, aa 241-400), and a C-terminal OSK domain (aa 401-606) **(Figure 4A).** Both the eLOTUS and the OSK domain bind to RNA *in vitro* (Ding *et al*, 2020; Jeske *et al*, 2015; Yang *et al*, 2015). In addition, the eLOTUS domain serves as a regulatory domain for the Vasa ATPase activity (Jeske *et al*, 2017). In Y2H experiments, Smaug bound to the DR fragment of Oskar **(Figure 4D).** Short Oskar that carried a deletion of the DR was not able to interact with the NTD of Smaug **(Figure 4E).** The DR of Oskar carries several regions with high sequence conservation across Drosophilids (**Supplemental Figure 7C**). Of these, Oskar 292-352 was the Smaug-binging region (SBR), as it was necessary and sufficient for Smaug interaction in Y2H assays **(Figures 4E).** Moreover, a shorter version of the Oskar SBR (aa 292-325), which showed highest conservation (**Supplemental Figure 7C**), fused to maltose-binding protein (MBP) was able to copurify the NTD of Smaug (aa 37-281) in an MBP pull-down assay **(Figure 4F).** Like Smoothened, Oskar bound to the NTD of Smaug, but not to the NTD of SAMD4A **(Supplemental Figure 7D)**.

### Predicted structure of the SBR of Oskar bound to the NTD of Smaug

Surface residues of the Smoothened-binding groove of the NTD of Smaug are relatively conserved across animals **(Supplemental Figure 2),** and we asked if the SBR of Short Oskar might bind to this groove too. We ran an AlphaFold2-multimer structural prediction (Evans *et al*, 2021; Jumper *et al*, 2021; Mirdita *et al*, 2022) using two copies each of the Smaug 70-280 (NTD) and the Oskar aa 295-300 as input sequences. We obtained a model of a heterotetramer, in which two α–helices covering part of the SBR of Oskar (aa 292-309) were placed into the Smoothened-binding groove of each D-PHAT domain of Smaug **(Figures 5A and 5B**). This shorter part of the SBR contains several conserved and highly conserved residues that contact the NTD of Smaug (**Figure 5C**). Importantly, using ReLo and Y2H assays, we found that the interaction between Short Oskar and Smaug is prevented when Oskar lacked the longer or shorter version of the SBR or when Smaug carried the point mutations in the NTD that also disrupted its interaction with Smoothened **(Figures 5D, 5E, and 5F**). Together, these data suggested that Smaug binds to Smoothened or Short Oskar in a similar fashion, and revealed the NTD of Smaug as peptide-binding domain.

**Figure 5.**
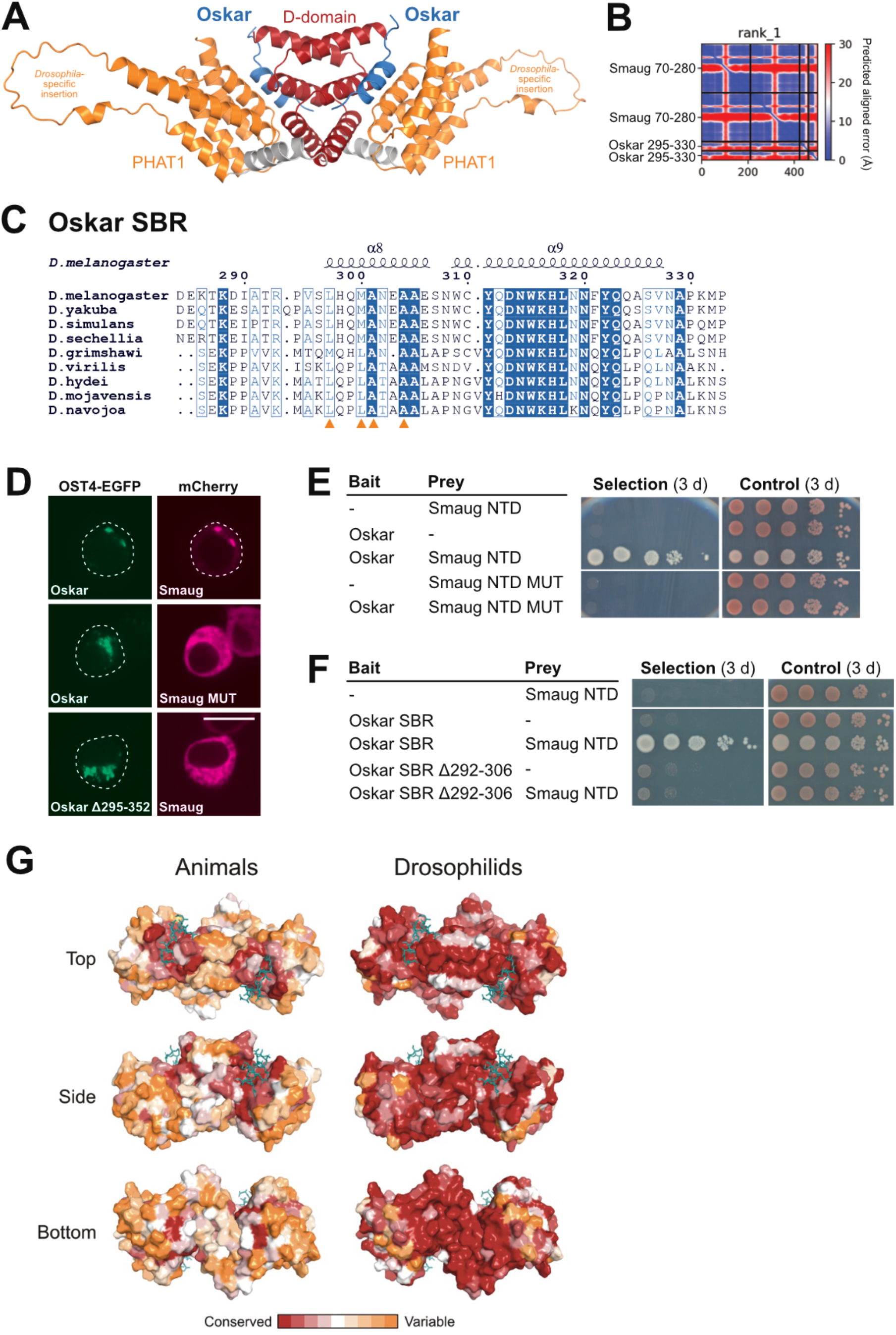
Predicted structure of Smaug-NTD in complex with the Oskar SBR. (A) Structural model of the Smaug-NTD in complex with the Oskar-SBR (blue) generated using AlphaFold2-multimer-version3 (Evans *et al*, 2021; Mirdita *et al*, 2022; Jumper *et al*, 2021). The model contains the *Drosophila*-specific insertion (aa 156-196) of the Smaug-NTD. (B) Predicted aligned error plot generated from the prediction run, showing a low positional error of the N-terminal part of the Oskar-SBR relative to the Smaug-NTD. (C) Sequence alignment of Oskar covering the SBR. Orange triangles indicate Oskar residues that are critical for Smaug binding based on the predicted structural model. (D) OST4-EGFP and mCherry fusions to the proteins indicated were coexpressed in S2R+ cells and their localization was analyzed by microscopy. The Smaug MUT (S250E, L253E) and the deletion of the SBR of Oskar (295-352) prevented the Smaug-Oskar interaction. Scale bar is 10 μm. (E, F) Yeast two-hybrid assays showing that the Smaug NTD MUT (S250E, L253E) or the deletion of aa 292-306 from the SBR (292-352) of Oskar prevented the Smaug-Oskar interaction. Selection medium lacked adenine and histidine. (G) Surface representation of the Smaug NTD colored according to residue conservation (Ashkenazy *et al*, 2016) considering sequences of Smaug orthologs from various animals (left panels) or Drosophilids only (right panels).

## DISCUSSION

Here, we present the crystal structures of the previously uncharacterized NTDs of human SAMD4A and *Drosophila* Smaug, which are composed of a D- and a PHAT-subdomain. These structural data revealed that Smaug and its animal orthologs contain not one but two PHAT domains, one connected to the dimerizing D-domain and one connected to the RNA-binding SAM domain. The D-domain has been predicted previously for Smaug and Vts1, and dimerization of the Vts1 D-domain was demonstrated (Tang *et al*, 2007). Furthermore, our structural analysis of the Smaug-Smoothened complex revealed that the D-domain not only functions as dimerization domain, but also forms a peptide binding groove together with the PHAT-subdomain.

Our data also demonstrated an interaction between the NTD of Smaug and a short part of the predominantly disordered region of Oskar. Structural modelling revealed a binding mode for the Oskar-Smaug interaction that is similar to the Smoothened-Smaug complex. The formation of such a protein binding groove is probably specific to the animal members of the Smaug protein family, as yeast Vts1 contains the D and the SAM domains, but lacks PHAT domains. The high conservation of the protein-binding groove of the NTD of metazoan Smaug/SAMD4 proteins suggests that SAMD4A, which does not bind to Oskar or Smoothened, may bind to a yet unknown partner in a similar fashion (**Figure 5G**).

Previously, a direct Oskar-Smaug interaction has been suggested and mapped to the SAM-PHAT2 domain of Smaug using classical Y2H assays (Dahanukar *et al*, 1999). The data from this interaction mapping raised the idea that Oskar might antagonize the Smaug repressive function by competing with the *nanos* SREs for binding to the RNA-binding SAM domain of Smaug (Dahanukar *et al*, 1999). However, the same lab also created transgenic flies to test mutations in Smaug that might interfere with Oskar interaction, without affecting RNA-binding, and reported that these substitutions had no effect on embryonic patterning (Dean *et al*, 2002). Using the split-ubiquitin-based Y2H, we did not observe a specific interaction between the SAM-PHAT2 region of Smaug and Short Oskar: we observed a similar degree of cell grow when the SAM-PHAT construct was coexpressed with the control plasmid as compared to its coexpression with Short Oskar. Instead, we found that Oskar bound to the NTD of Smaug. Dahanukar et al. did not test a Smaug fragment comprising the full NTD, but only a truncated fragment (1-242) lacking a large part of the PHAT1 subdomain. This truncated Smaug construct did not show an Oskar interaction (Dahanukar *et al*, 1999). Our Y2H data, structural modeling and mutagenesis all lead to the conclusion that Oskar binds to the NTD of Smaug, but it remains unclear if the interaction contributes to a block of Smaug binding to *nanos* mRNA and if so, how. As Oskar did not bind to the RNA-binding SAM domain of Smaug, and the NTD of Smaug did not bind to RNA in *in vitro* binding experiments (**data not shown**), Oskar seems not to actively interfere with binding of Smaug to *nanos* mRNA. UV crosslink experiments indicate that Short Oskar binds to *nanos* mRNA *in vivo* (Jeske *et al*, 2015). However, from this experiment it is unclear whether Oskar is capable of recognizing *nanos* mRNA directly or whether the RNA-binding specificity is mediated by another RNA-binding protein *in vivo*, such as Smaug. The eLOTUS domain of Short Oskar binds to the DEAD-box RNA helicase Vasa and stimulates its ATPase activity (Jeske *et al*, 2017), suggesting a contributing function of Vasa in *nanos* mRNA control. Whether indeed Oskar works together with Vasa to prevent the Smaug-mediated repression of *nanos* mRNA at the posterior pole remains to be investigated.

Our biochemical and structural data demonstrate a direct interaction between Smoothened and Smaug, yet the function of this interaction is unclear. In cell culture assays, Smaug is phosphorylated upon HH signaling and this phosphorylation depends on Smoothened (Bruzzone *et al*, 2020). Further studies in cell culture, in wing imaginal discs, and fly wings suggested that Smaug phosphorylation leads to a reduced mRNA repressive activity (Bruzzone *et al*, 2020). Smaug has an established function as repressor of maternal mRNAs during early *Drosophila* embryogenesis. Consistent with this function, Smaug protein expression is highest between 0 and 3 hours of embryonic development (Dahanukar *et al*, 1999; Gramates *et al*, 2022; Casas-Vila *et al*, 2017) (**Supplemental Figure 8A)**. In contrast, Smaug· protein levels are at the detection limit in all developmental stages beyond the larvae stage L3 (Casas-Vila *et al*, 2017) (**Supplemental Figure 8A),** and Smaug defects were reported earlier to not cause any phenotype in the adult fly (Dahanukar *et al*, 1999).

Keeping these earlier observations in mind, we asked if there might be a function for the Smaug - Smoothened interaction in the early embryo. During early embryogenesis, the protein expression pattern of Smoothened is reciprocal to the one of Smaug: Smoothened protein levels are low in early embryos, and strongly enhanced after 3 h of development (Heuvel & Ingham, 1996; Casas-Vila *et al*, 2017) (**Supplemental Figure 8A**). This reciprocal expression pattern might indicate that one protein regulates the level of the other protein. Smaug protein is actively cleared during the MZT (after ca. 3 h), and its degradation is initiated by SCF-mediated ubiquitination (Cao *et al*, 2022, 2020). Furthermore, it was suggested that Smaug is recognized by the F-box proteins Slmb (the *Drosophila* β-TrCP ortholog) and Bard, with Bard exhibiting a predominant and a timing role in Smaug degradation (Cao *et al*, 2022, 2020). In our ReLo assays, we did not observe the Smaug - Slmb interaction, but detected the Smaug - Bard interaction (**Supplemental Figure 8B**). Often the recognition of target proteins by F-box proteins, such as Slmb, depends on prior phosphorylation of the target (Morais-de-Sá *et al*, 2013; Mason & Laman, 2020; Frescas & Pagano, 2008; Jia *et al*, 2005). Mouse and human SAMD4A/Smaug1 were reported to interact with 14-3-3 proteins (Chen *et al*, 2014b; Fehilly *et al*, 2023), a protein family known to recognize phosphorylated sequences within their protein partners (Obsilova & Obsil, 2022). We detected the *Drosophila* Smaug interaction to 14-3-3ζ and 14-3-3ε in our ReLo assays (**Supplemental Figure 8C**), suggesting that Smaug is phosphorylated in these cells. Furthermore, Smaug phosphorylation has been described in the embryo and in *Drosophila* Cl8 cell culture (Bruzzone *et al*, 2020; Zhai *et al*, 2008). In Cl8 cells, Smaug phosphorylation was dependent on both Smoothened expression and HH signaling. Additional data suggested that upon HH signaling, Smoothened binds simultaneously to Smaug and the serin/threonine kinase Fused, and thereby mediates the phosphorylation of Smaug by Fused (Bruzzone *et al*, 2020). Using our ReLo assays, we confirmed a direct interaction between Smoothened and Fused and, consistent with previous Y2H data (Malpel *et al*, 2007), we mapped Smoothened binding to the C-terminal regulatory domain of Fused (**Supplemental Figure 8D**). Considering this chain of interactions and activities, we are wondering whether a Smaug phosphorylation event mediated by Smoothened might be relevant for the SCF-mediated ubiquitination and degradation of Smaug in the early embryo. In this manner, Smoothened might contribute to the timing of the MZT in the early embryo, an exciting hypothesis that needs to be tested in the future.

## METHODS

### Cloning of DNA constructs

For the generation of the ReLo cloning vector pAc5.1-mEGFP (C-ter) (EB2), a unique FspAI restriction site was introduced 5’ of the mEGFP coding sequence of the pAc5.1-mEGFP vector (T6-MJ) The pAc5.1-OST4-EGFP vector (JK268) was generated by inserting the *Saccharomyces cerevisiae* OST4 protein sequence amplified from pDHB1 (DualSystems Biotech) into the pAc5.1-EGFP plasmid (code) using the KpnI restriction site. In the next step, the unique FspAI restriction site was introduced 3’ to to the EGFP coding sequence. The FspAI sites were used to insert PCR-amplified inserts with blunt-end cloning. All other ReLo cloning vectors were described previously (Salgania *et al*, 2022; Jeske *et al*, 2017; Behm-Ansmant *et al*, 2006). The Y2H cloning vectors were described previously (Pekovic *et al*, 2022; Jeske *et al*, 2015).

The pMJ-His *E. coli* expression vector was generated by introducing a unique ScaI blunt end restriction site followed by a stop codon as close as possible downstream of the TEV protease recognition site of the pET-M11 vector. The pMJ-His-MBP vector was created by modification of the pET-M44 vector in a similar manner. The pMJ-GST vector was generated from the pGEX-6P-1 plasmid by introducing a unique SmaI blunt end restriction site as close as possible downstream to the HRV 3C protease recognition sequence.

To generate the fusion construct for crystallization, the sequence of Smaug 73-278Δ156-196 was amplified from the pMJ-His-Smaug 70-281Δ156-196 plasmid (AG4) and ligated into the pMJ-His vector (T43-MJ) digested with ScaI. The sequence of Smo 970-1003 E975Q-(GGS)_4_ was amplified from a synthetic DNA fragment codon-optimized for *E. coli* (gBlock from IDT) and ligated into the pMJ-His-Smaug 73-278Δ156-196 vector, which was opened by PCR amplification 5’ of the Smaug 73-278Δ156-196 sequence. Smo 970-1003 E975Q was deleted from the obtained plasmid by PCR amplification and Smo 970-1003 amplified from pAc 5.1-mEGFP-Dm Smo 556-1035 (JK168) was inserted by blunt-end ligation.

Detailed information on all plasmids used in this study is provided in **Supplementary Table 2**.

### Protein purification

Proteins were expressed using the autoinduction method (Studier, 2005). Briefly, Rosetta™ 2 competent cells (Novagen) were transformed with the protein expression vector and grown at 37°C in LB medium supplemented with antibiotics. The following day, TB medium supplemented with antibiotics was inoculated with the overnight culture and grown at 22°C or 23°C for 27 h. For SeMet labeling of the *Hs*-SAMD4A 2-156 protein, the PASM-5052 expression medium (Studier, 2005) was used. Proteins were purified using affinity chromatography (Ni2+-NTA or glutathione agarose), followed by ion exchange chromatography and size exclusion chromatography. Unless indicated otherwise, proteins were eluted in a buffer containing 20 mM Tris-Cl, pH 7.5 and 150 mM NaCl. If needed, the His tag was removed by TEV protease cleavage prior to the ion exchange chromatography step.

### Protein-protein interaction assays

ReLo assays (Salgania *et al*, 2022), Y2H assays (Jeske *et al*, 2015), and GST pulldown assays (Jeske *et al*, 2017) were performed essentially as described. MBP pulldown assays were performed similar to GST pull down assays using Amylose resin (New England Biolabs) and 1 nmol of MBP-Oskar 292-325 and 5 nmol of Smaug 37-281.

Isothermal titration calorimetry (ITC) was performed at 15°C using a MicroCal PEAQ instrument (Malvern Panalytical Ltd) and a buffer containing 20 mM Tris-Cl, pH 7.5 and 150 mM NaCl. The syringe contained 200 μM of Smaug 70-281 protein, which was titrated in 19 injections into the cell containing 20 μM of a Smoothened 970-1003 synthetic peptide (Peptide Specialty Laboratories GmbH, Germany). The first injection consisted of 0.4 μl, the consecutive injections had 2 μl. Reference power was set to 10 μW, the stir speed was 750 rpm. In the control experiment, the buffer was titrated into the cell with the peptide. The results were fitted and analyzed with the MicroCal PEAQ-ITC Analysis Software.

Multiangle-light-scattering (MALS) experiments were performed using an ÅKTA purifier system (Cytiva) equipped with a Superdex 200 Increase 10/300 column (Cytiva) and connected to the MALS detector DAWN 8+ (Wyatt Technology) and SEC-3010 RI Detector (WGE Dr Bures GmbH, Germany). The system and the proteins were in buffer containing 20 mM Tris-Cl, pH 7.5 and 150 mM NaCl. 100 μl of Smaug 70-281 (508 μM) or Hs-SAMD4A 2-156 (1.1 mM) each supplemented with 20 mM DTT was injected. The data were fitted and analyzed with the Astra 6 software (Wyatt Technology).

### Crystallization and structure determination

The crystallization of His-Hs-SAMD4A 2-156 was performed using the hanging drop vapor diffusion method. 2 μl of native His-SAMD4A 2-156 (23 mg/ml) or SeMet-labelled His-SAMD4A 2-156 (37.5 mg/ml) in crystallization buffer (20 mM Tris-Cl, pH 7.5 and 150 mM NaCl, supplemented with 5 mM DTT) were mixed with 2 μl precipitant (0.1 M sodium citrate, pH 5.6 and 8% jeffamine M-600, pH 7) and incubated at 18°C. Crystals appeared after one day and were cryogenically protected by supplementing them with 35% or 40% (v/v) glycerol prior to flash-freezing in liquid nitrogen. Diffraction data for native and SeMet crystals were collected at P14 using 0.9686 Å and at P13 beamline using 0.9800 Å, respectively, operated by EMBL Hamburg at the PETRA III storage ring (DESY, Hamburg, Germany). The diffraction data were processed anisotropically using STARANISO (Global Phasing Limited). The phases were determined with multi-wavelength anomalous dispersion using the reflections from the peak of SeMet derivatives and the native crystal using the AutoSol program of the PHENIX suite (Adams *et al*, 2010).

The Smoothened 970-1003-(GGS)_4_-Smaug 73-278Δ156-196 fusion protein at 11.6 mg/mL concentration (supplemented with 5 mM DTT) was crystallized by mixing with an equal volume of precipitant (0.1 M HEPES pH 7.5, 5% (w/v) PEG3000, 20% (w/v) PEG400) using the sitting-drop vapor diffusion method at 18°C. Rod/needle-shaped single crystals appeared after seven days. The crystals were cryogenically protected by supplementing them with 10% glycerol prior to flash-freezing in liquid nitrogen. The diffraction data were processed anisotropically using STARANISO (Global Phasing Limited) and the structure was solved by molecular replacement using the structure of the SAMD4A-NTD dimer as a search template. Structures were refined using phenix.refine and Coot (Afonine *et al*, 2012; Emsley & Cowtan, 2004), and figures were generated using PyMol. All data collection and refinement statistics are provided in **Supplementary Table 1**.

## ACKNOWLEDGEMENTS

We thank Jochen Baßler, Anastasija Gricenko, Felia Haffelder, David Hauser, Maik Schauerte, Alina Schneider, Jonas Weidenhausen, Julan Weng, and Janka Zsok for technical support. We thank Elisa Izaurralde for plasmids. We thank Elmar Wahle and Craig Smibert for constructive feedback on the manuscript. We thank the Nikon Imaging Center at the University of Heidelberg for access to microscopes and the BZH Protein Crystallization Platform. We gratefully acknowledge the P13 and P14 beamlines operated by EMBL Hamburg at the PETRA III storage ring (DESY, Hamburg, Germany) and the ID-30B beamline at the ESRF (Grenoble, France). We would like to thank Isabel Bento (DESY) and Nicolas Coquelle (ESRF) for the assistance in using the beamlines. We gratefully acknowledge the data storage service SDS@hd supported by the Ministry of Science, Research and the Arts Baden-Württemberg (MWK) and the German Research Foundation (DFG) through the grants INST 35/1314-1 FUGG and INST 35/1503-1 FUGG. This work was funded by the Emmy-Noether Program of the German Research Foundation (DFG; JE-827/1-1).

## ACCESSION NUMBERS

The structures reported here have been deposited to the Protein Data Bank (PDB) and are available under accession numbers **xxxx** (SAMD4A-NTD) and **xxxx** (Smaug-NTD - Smoothened peptide complex).

