## Supplemental data for "Structural basis for binding of Smaug to the GPCR Smoothened and to the germline inducer Oskar"

Inventory:

Supplemental Figure 1

Supplemental Figure 2

Supplemental Figure 3

Supplemental Figure 4

Supplemental Figure 5

Supplemental Figure 6

Supplemental Figure 7

Supplemental Figure 8

Supplemental Table 1

Supplemental Table 2

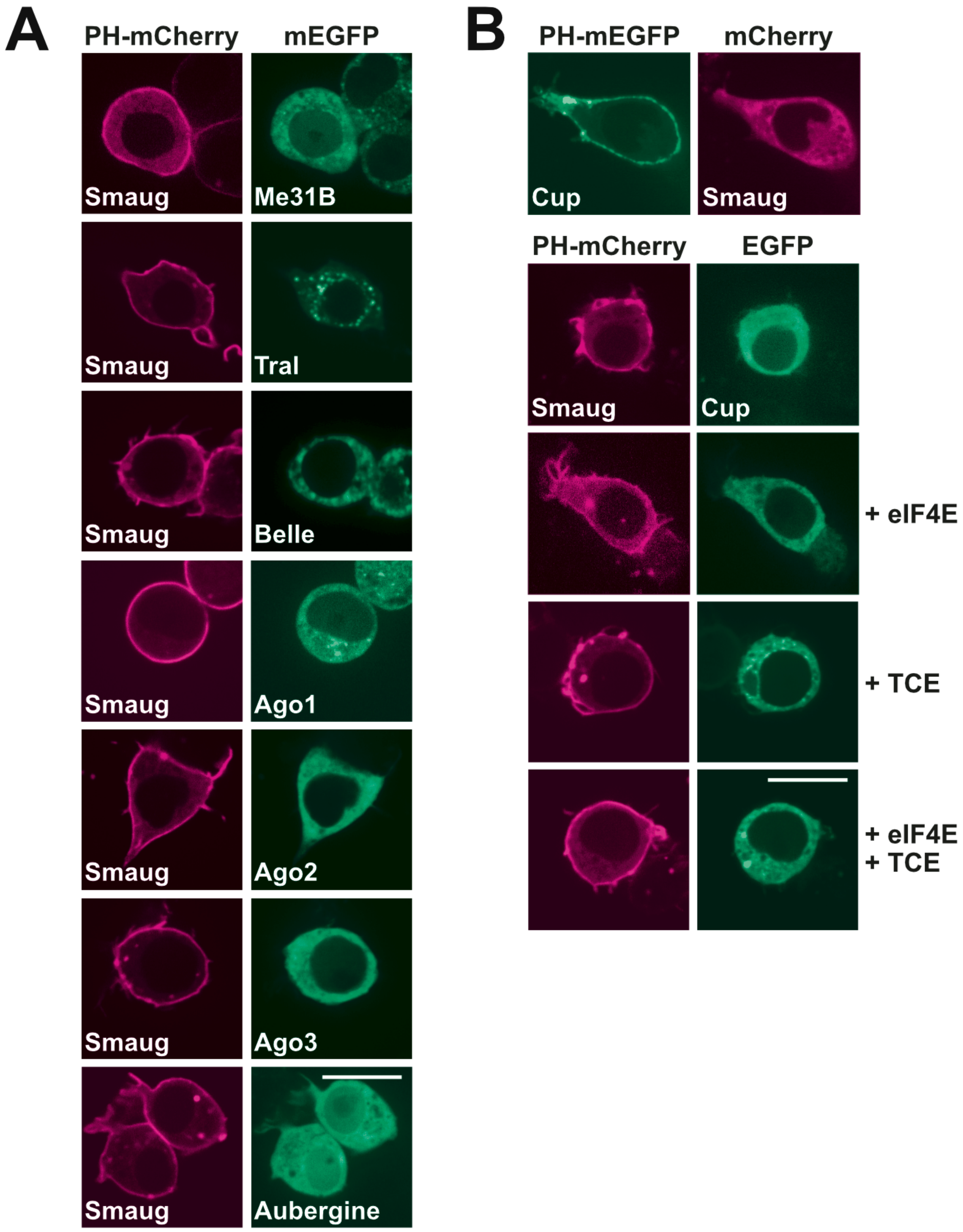

**Supplemental Figure 1. Smaug PPI testing using ReLo assays.**
(A) PH-mCherry-Smaug and mEGFP fusions to the proteins indicated were coexpressed in S2R+ cells and their localization was analyzed by microscopy. Aubergine was fused to EGFP. Proteins indicated did not interact with Smaug in the ReLo assay, as concluded by their lack of membrane relocalization. (B) Fusion proteins indicated were coexpressed in S2R+ cells and their localization was analyzed by microscopy. In the lower three panels PH-mCherry-Smaug and EGFP-Cup were coexpressed together with nonfluorescent eIF4E protein and/or a luciferase construct containing the *nanos* 3'UTR translation control element (TCE). In no instance, Smaug and Cup showed an interaction. Scale bars are 10 µm.

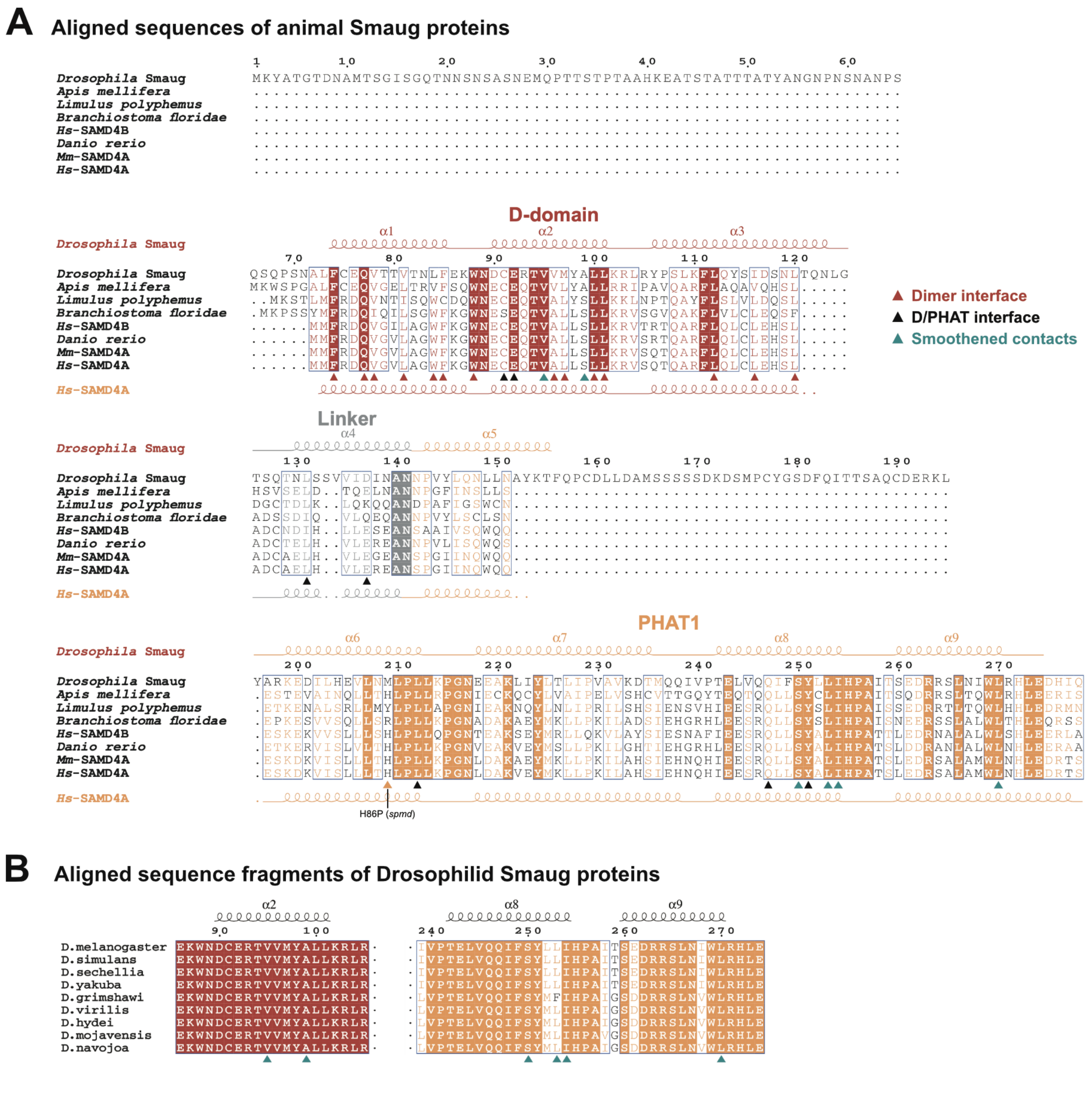

**Supplemental Figure 2. Smaug sequence conservation.**
(A) Multiple sequence alignment of animal Smaug family members of the species indicated generated with MUSCLE (Edgar, 2004) and visualized with ESPript 3 (Robert & Gouet, 2014). Arrowheads indicate residues involved in dimerization (red), the D-PHAT interaction (black), or Smoothened binding (deepteal). (B) Multiple sequence alignment of *Drosophila* Smaug proteins.

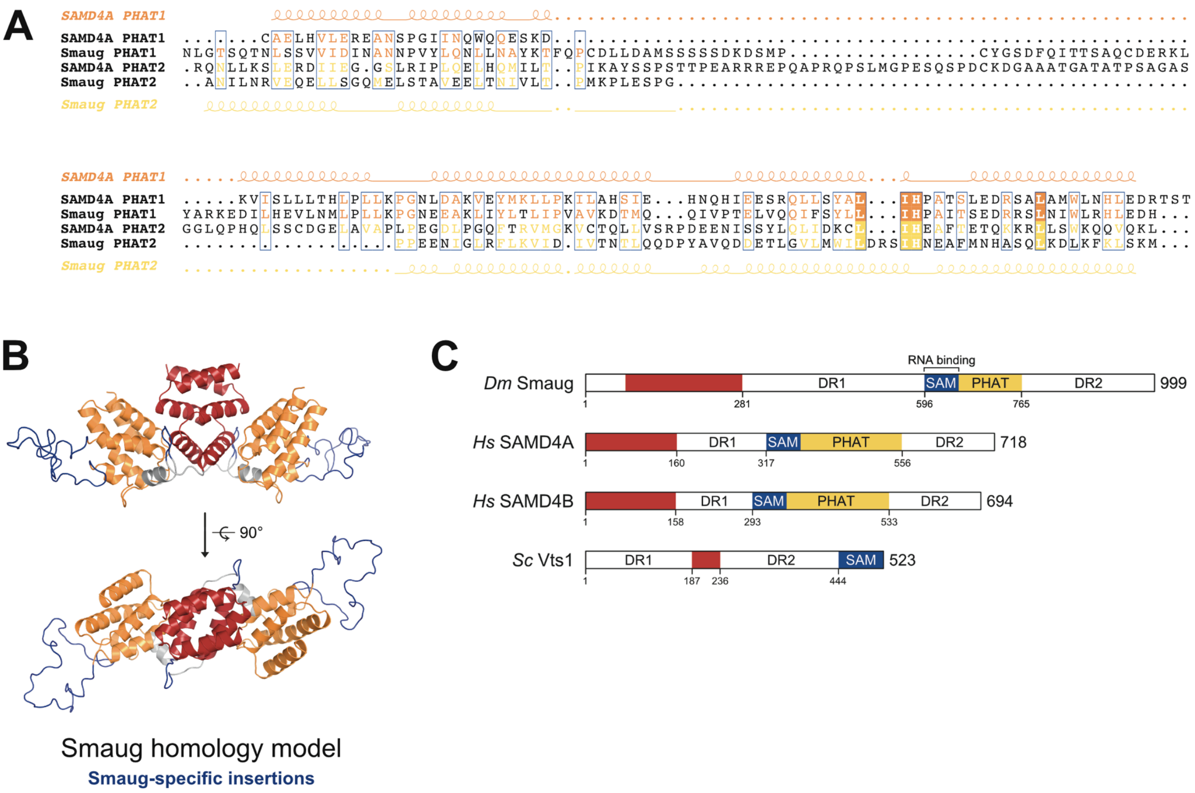

**Supplemental Figure 3. PHAT domains in the Smaug protein family.**
(A) Sequence alignment generated with MUSCLE (Edgar, 2004) and visualized with ESPript 3 (Robert & Gouet, 2014) using the PHAT1 and PHAT2 domain sequences of Smaug and SAMD4A. Smaug PHAT1 and SAMD4A PHAT2 domains contain longer insertions. (B) Homology model of the Smaug NTD based on the SAMD4A-NTD created with SWISS-MODEL (Waterhouse *et al*, 2018). Specific protein insertions are indicated in dark blue color. (C) Protein domain organization of the proteins indicated. Yeast Vts1 lacks PHAT domains. *Dm*, *Drosophila melanogaster*; *Hs*, *Homo sapiens*; *Sc*, *Saccharomyces cerevisiae*.

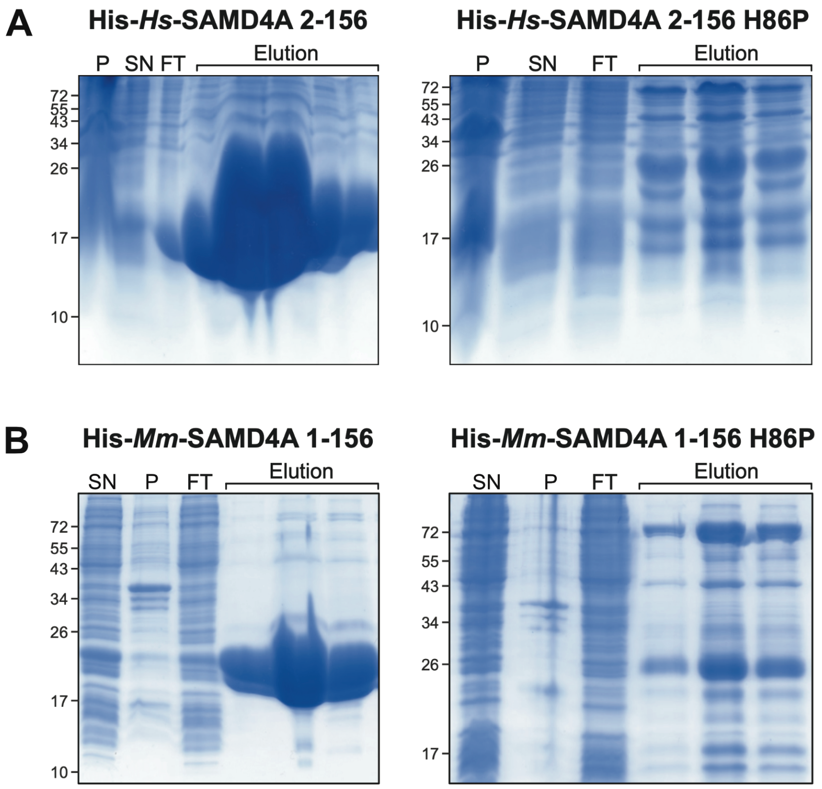

**Supplemental Figure 4. The *spmd* mutation H86P caused insolubility of the NTD of human and mouse SAMD4A.**

His-tagged constructs as indicated were recombinantly expressed and purified in parallel from *E. coli*. Pellet (P), supernatant (SN), flowthrough (FT) and elution fractions obtained during Ni-NTA affinity purification were loaded on an SDS-PAGE and analyzed by Coomassie staining. Protein size markers are in kDa. (A) Human SAMD4A. (B) Mouse SAMD4A.

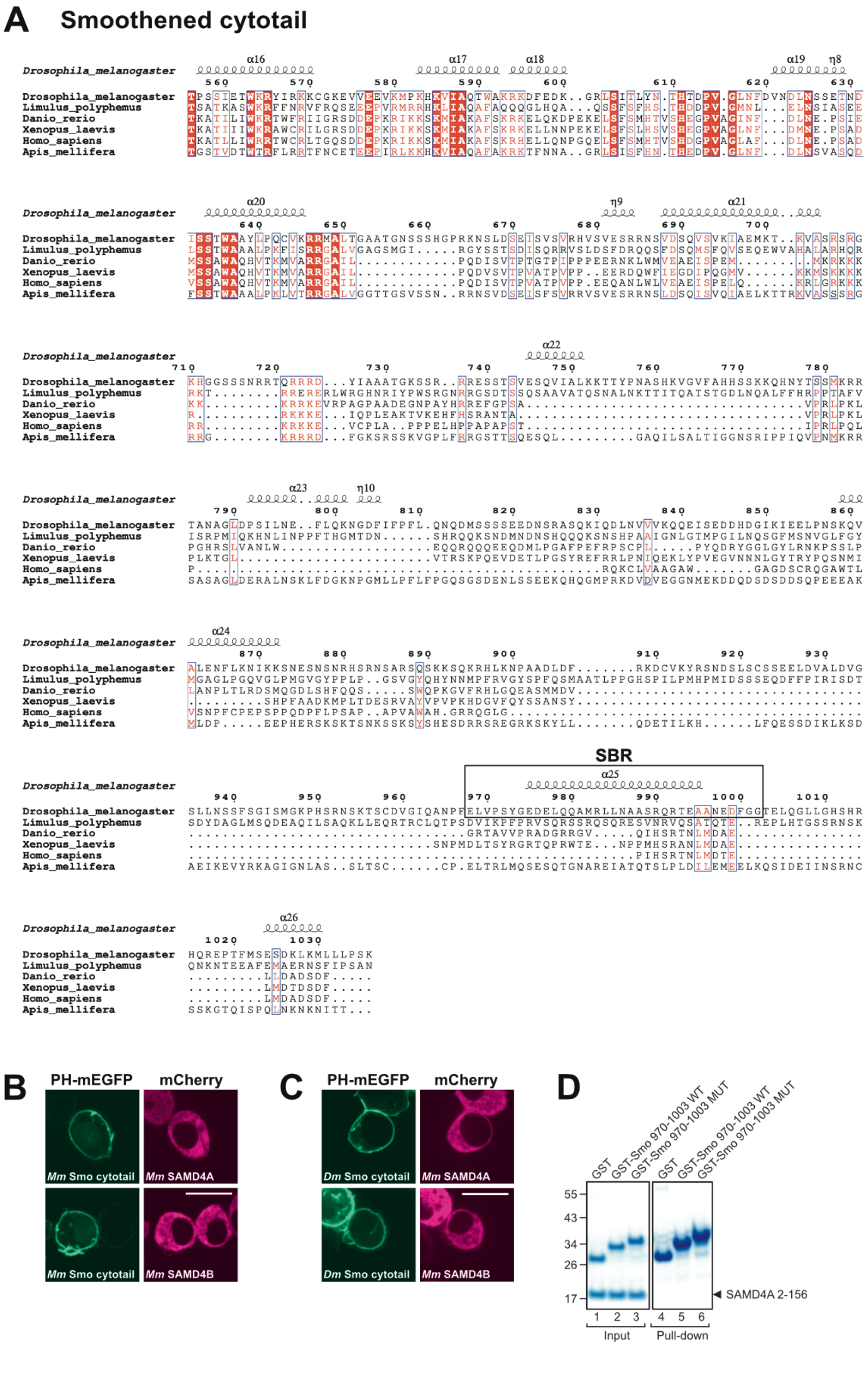

**Supplemental Figure 5. The Smaug - Smoothened interaction is specific to *Drosophila*.**
(A) Multiple sequence alignment of the Smoothened Cytotail using various animal sequences created with MUSCLE (Edgar, 2004) and visualized with ESPript 3 (Robert & Gouet, 2014). (B, C) PH-mEGFP and mCherry fusions to the proteins indicated were coexpressed in S2R+ cells and their localization was analyzed by microscopy. The Cytotail of mouse (Mm) Smoothened (B) or *Drosophila* (Dm) Smoothened (C) did not interact with mouse SAMD4A or SAMD4B in the ReLo assay. Scale bars are 10 µm. (D) GST pull-down assay showing that the *Drosophila* Smoothened SBR did not bind to the NTD of human SAMD4A. The experiment was performed in parallel with the one shown in main **Figure 3E**. Protein markers are in kDa.

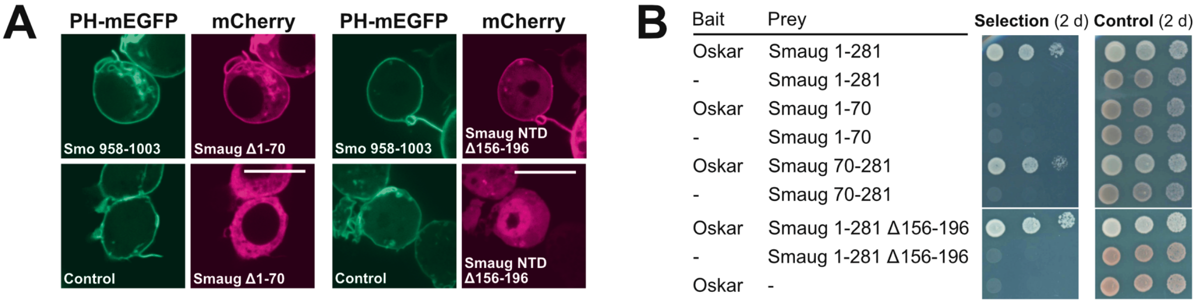

**Supplemental Figure 6. Smaug NTD extensions and insertions were not necessary for binding to Smoothened or Oskar.**
(A) PH-mEGFP and mCherry fusions to the proteins indicated were coexpressed in S2R+ cells and their localization was analyzed by microscopy. Scale bars are 10 μm.
(B) Split-ubiquitin yeast two-hybrid assay with bait and prey vectors coexpressed as indicated. Cells were spotted in three ten-fold dilutions and grown for two days. Selection medium lacked adenine and histidine.

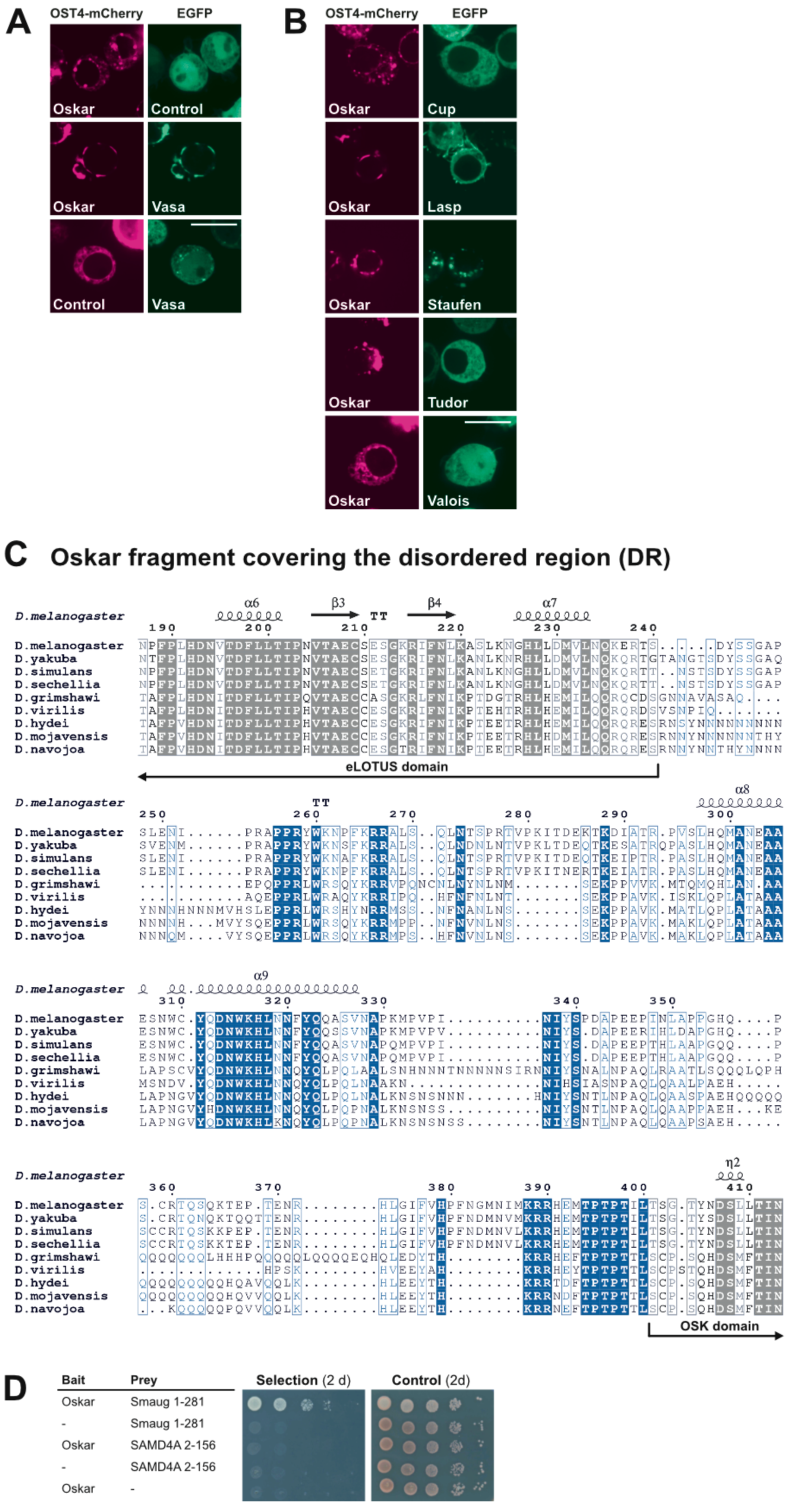

**Supplemental Figure 7. Short Oskar PPI testing.**
(A, B) EGFP and OST4-mCherry fusions to the proteins indicated were coexpressed in S2R+ cells and their localization was analyzed by microscopy. Control indicates the mEGFP vector lacking a fusion. Short Oskar interacted with Vasa (A) but not with other candidates (B) in the ReLo assay. Scale bars are 10 µm. (C) Multiple sequence alignment of an Oskar fragment covering the disordered region (DR) created with MUSCLE (Edgar, 2004) and visualized with ESPript 3 (Robert & Gouet, 2014). (D) Split-ubiquitin yeast two-hybrid assay as performed as described in **Supplemental Figure 6B**.

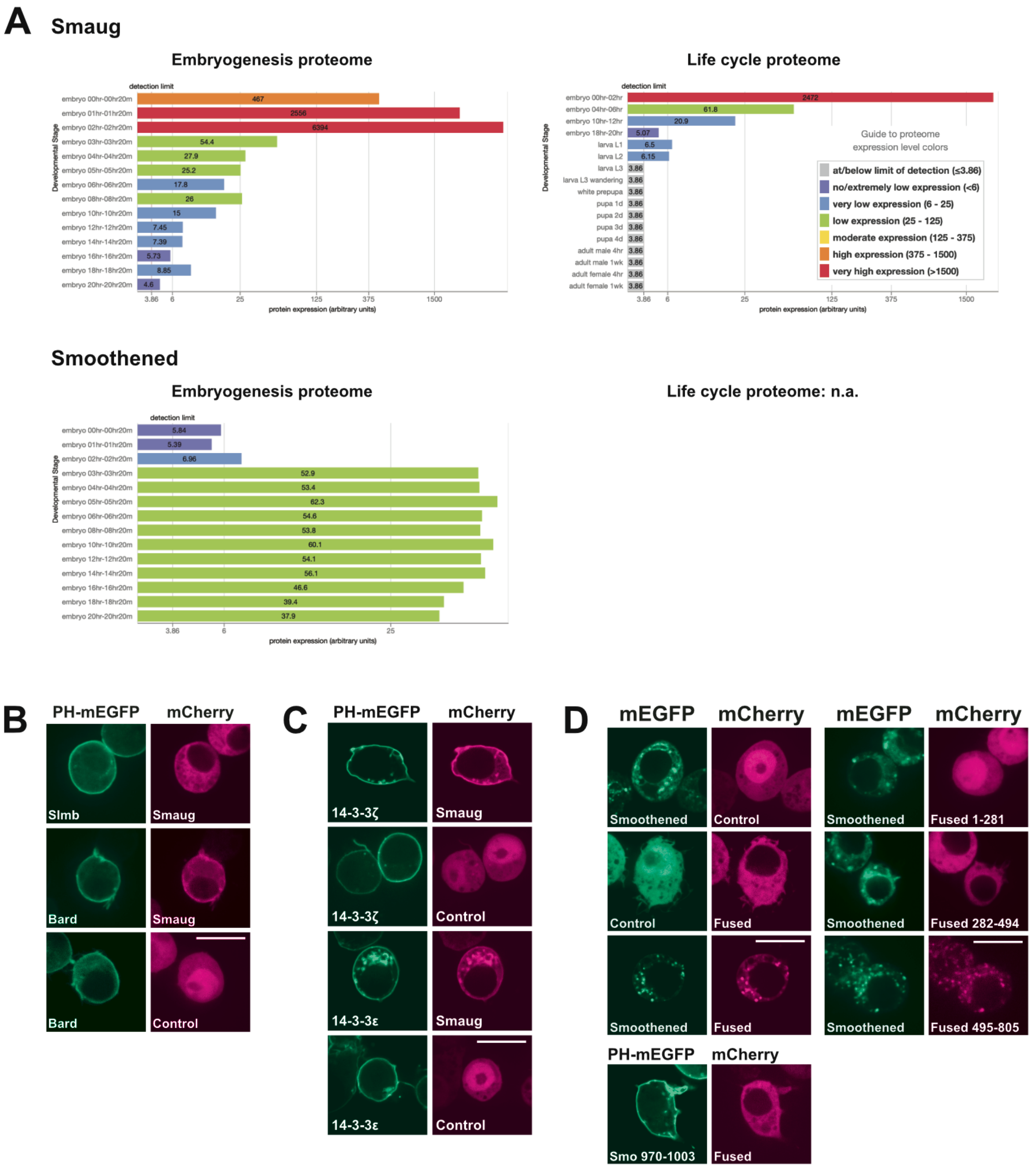

**Supplemental Figure 8. Protein expression data and additional interactions of Smaug and Smoothened.**

(A) Smaug and Smoothened protein expression data during *Drosophila* embryogenesis or the life cycle (Casas-Vila *et al*, 2017; Gramates *et al*, 2022). In 0-2 h and 20 min early embryos, Smaug protein levels are very high and Smoothened levels are at the detection limit. After 3 h of embryogenesis, Smoothened expression is low but present, while Smaug levels are strongly reduced. Smaug protein levels are very low in larva and at the detection limit in late larvae, pupa, and adult tissue. (B-D) PH-mEGFP, EGFP, and mCherry fusions to the proteins indicated were coexpressed in S2R+ cells and their localization was analyzed by microscopy. Control indicates the mCherry vector lacking a fusion. Smoothened carried a C-terminal mEGFP tag to maintain its membrane localization. Smaug relocalized with Bard but not with Slmb (B), and with 14-3-3ζ and 14-3-3ε (C); Smoothened bound to the C-terminal domain (495-805) of Fused (D) in the ReLo assay. Scale bars are 10 µm.

**Supplemental Table 1. Data collection and refinement statistics.**
Statistics for the highest-resolution shell are shown in parentheses.

| | **SAMD4A** | SAMD4A SeMet (x5_peak) | SAMD4A SeMet (x6_peak) | SAMD4A SeMet (x10_peak) | **Smg-Smo** |
|---|---|---|---|---|---|
| Wavelength (Å) | 0.968 | 0.980 | 0.980 | 0.980 | 0.976 |
| Beamline | P14 | P13 | P13 | P13 | ID-30B |
| Resolution range (Å) | 63.05 - 1.62 (1.7 - 1.62) | 71.54-3.0 (3.19-3.0) | 71.54-3.0 (3.19-3.0) | 71.54-3.0 (3.19-3.0) | 66.33- 2.00 (2.23 - 2.00) |
| Space group | C 2 2 21 | C 2 2 21 | C 2 2 21 | C 2 2 21 | P 61 2 2 |
| Unit cell (a b c a b g) | 81.8 142.8 137.1 90 90 90 | 81.9 142.8 137.1 90 90 90 | 81.6 142.8 137.1 90 90 90 | 81.9 142.8 137.1 90 90 90 | 76.6 76.6 268.5 90 90 120 |
| Total reflections | 1238981 (64282) | 218325 (37049) | 217266 (37681) | 220139 (37688) | 216855 (12281) |
| Unique reflections | 89788 (4488) | 31152 (5247) | 31127 (5228) | 31099 (5197) | 21810 (1090) |
| Multiplicity | 13.8 (14.3) | 7.0 (7.1) | 7.0 (7.2) | 7.1 (7.3) | 9.9 (11.3) |
| Completeness (spherical, %) | 87.5 (29.5) | 15.0 (64.3) | 15.0 (64.1) | 15.0 (63.7) | 66.8 (12.5) |
| Completeness (ellipsoidal, %) | 95.2 (60.2) | - | - | - | 94.0 (78.4) |
| Mean I/sigma(I) | 19.2 (0.4) | 27.18 (13.26) | 31.50 (13.87) | 26.28 (10.10) | 7.0 (0.9) |
| R-merge | 0.060 (4.38) | | | | 0.196 (2.36) |
| R-meas | 0.062 (4.55) | 0.065 (0.157) | 0.052 (0.137) | 0.072 (0.230) | 0.206 (2.47) |
| R-pim | 0.017 (1.20) | | | | 0.063 (0.72) |
| CC1/2 | 0.999 (0.34) | 0.999 (0.992) | 0.999 (0.993) | 0.999 (0.986) | 0.996 (0.70) |
| Reflections used in refinement | 89724 | | | | 21804 |
| Reflections used for R-free | 2001 | | | | 1998 |
| R-work (%) | 20.48 | | | | 27.18 |
| R-free (%) | 22.15 | | | | 29.14 |
| CC (work) | 0.951 | | | | 0.902 |
| CC (free) | 0.946 | | | | 0.868 |
| Number of non-hydrogen atoms | 4253 | | | | 2950 |
| macromolecules | 3862 | | | | 2838 |
| ligands | 25 | | | | 37 |
| solvent | 366 | | | | 97 |
| Protein residues | 479 | | | | 350 |
| RMS (bonds) | 0.008 | | | | 0.012 |
| RMS (angles) | 1.18 | | | | 1.54 |
| Ramachandran favored (%) | 98.94 | | | | 97.95 |
| Ramachandran allowed (%) | 1.06 | | | | 2.05 |
| Ramachandran outliers (%) | 0.00 | | | | 0.00 |
| Rotamer outliers (%) | 0.47 | | | | 0.61 |
| Clashscore | 6.91 | | | | 14.87 |
| Average B-factor | 41.87 | | | | 40.67 |
| macromolecules | 41.42 | | | | 40.71 |
| ligands | 51.12 | | | | 32.57 |
| solvent | 45.95 | | | | 40.70 |
| PDB code | xxxx | | | | xxxx |

**Supplemental Table 2. DNA constructs used in this study.**
All constructs are *Drosophila melanogaster* sequences, if not indicated otherwise. Plasmids pDHB1 (T9-MJ) and pPR3-N (T11-MJ) are from DualSystems Biotech, Zurich, Switzerland. Plasmids pDHB1-Short Oskar (139-606) (L2-MJ), pDHB1-Oskar 139-240 (LOTUS) (L4-MJ), pDHB1-Oskar 241-400 (DR) (L7-MJ), and pDHB1-Oskar 401-606 (OSK) (L8-MJ) have been described in (Jeske *et al*, 2015). Plasmids pAc5.1-mEGFP (T6-MJ), pAc5.1-mCherry (T7-MJ), pAc5.1-EGFP (T5-MJ), and pAc5.1-EGFP-Vasa (F15-MJ) have been described in (Jeske *et al*, 2017). Plasmids pAc5.1-PH-mEGFP (JM50), pAc5.1-PH-mEGFP-Cup (JM51), pAc5.1-PH-mCherry (HK49), pAc5.1-OST4-mCherry (XH26), pAc5.1-OST4-mCherry-Short Oskar (139-606) (H3-MJ), pAc5.1-EGFP-Aubergine (F20-MJ), and pAc5.1-EGFP-Cup (F23-MJ) have been described in (Salgania *et al*, 2022). Plasmids pDHB1-MJ (JK16), pPR3-N-MJ (JK18), and pAc5.1-EGFP-Smaug (F31-MJ) have been described in (Pekovic *et al*, 2022).

| **Vector** (insertion site) (code) | **Final DNA construct** | **DNA template information** | **Code** |
|---|---|---|---|
| **pDHB1** (bait) (NcoI) (T9-MJ) | pDHB1-**Short Oskar** (139-606) | (Jeske *et al*, 2015) | L2-MJ |
| | pDHB1-**Oskar 139-240 (LOTUS)** | (Jeske *et al*, 2015) | L4-MJ |
| | pDHB1-**Oskar 241-400 (DR)** | (Jeske *et al*, 2015) | L7-MJ |
| | pDHB1-**Oskar 401-606 (OSK)** | (Jeske *et al*, 2015) | L8-MJ |
| | pDHB1-**Oskar 292-352 (SBR)** | pPR3-N-Oskar (N1-MJ) | JK35 |
| **pDHB1-MJ** (bait) (Eco47III) (JK16) (Pekovic *et al*, 2022) | pDHB1-MJ-**Oskar Δ241-400** | pDHB1-Short Oskar (L2-MJ) | FH10 |
| | pDHB1-MJ-**Oskar Δ292-352** | pDHB1-Short Oskar (L2-MJ) | JM44 |
| | pDHB1-MJ-**Oskar 292-352 (SBR) Δ292-306** | pDHB1-Short Oskar (L2-MJ) | JM43 |
| **pPR3-N** (prey) (BamHI/EcoRI) (T11-MJ) | pPR3-N-**Smaug** | *smaug* cDNA | N16-MJ |
| **pPR3-N-MJ** (prey) (SmaI) (JK18) (Pekovic *et al*, 2022) | pPR3-N-MJ-**Smaug 1-281 (NTD)** | *smaug* cDNA | JK98 |
| | pPR3-N-MJ-**Smaug 1-281 S250E/L253E (NTD MUT)** | pAc5.1-PH-mCherry-Smaug 1-281 (S250E/L253E) (JM265) | JM271 |
| | pPR3-N-MJ-**Smaug 282-595 (DR1)** | *smaug* cDNA | JK100 |
| | pPR3-N-MJ-**Smaug 596-764 (SAM+PHAT)** | *smaug* cDNA | JK102 |
| | pPR3-N-MJ-**Smaug 765-999 (DR2)** | *smaug* cDNA | JK103 |
| | pPR3-N-MJ-**Smaug 1-70** | *smaug* cDNA | FH26 |
| | pPR3-N-MJ-**Smaug 70-281** | *smaug* cDNA | JK149 |

| | | | |
|---|---|---|---|
| | pPR3-N-MJ-**Smaug 1-281 Δ156-196** | pPR3-N-MJ-Smaug 1-281 (JK98) | AG6 |
| | pPR3-N-MJ-**human SAMD4A 2-156** | synthetic DNA sequence codon-optimized for *E. coli* (IDT) | FH19 |
| **pAc5.1-EGFP** (N-ter) (EcoRV) (T5-MJ) (Jeske *et al*, 2017) | pAc5.1-EGFP-**Smaug** | (Pekovic *et al*, 2022) | F31-MJ |
| | pAc5.1-EGFP-**Aubergine** | (Salgania *et al*, 2022) | F20-MJ |
| | pAc5.1-EGFP-**Vasa** | (Jeske *et al*, 2017) | F15-MJ |
| | pAc5.1-EGFP-**Cup** | (Salgania *et al*, 2022) | F23-MJ |
| | pAc5.1-EGFP-**Lasp** | cDNA | F24-MJ |
| | pAc5.1-EGFP-**Staufen** | cDNA | F32-MJ |
| | pAc5.1-EGFP-**Tudor** | Assembly of 3 *tudor* cDNA fragments | F38-MJ |
| | pAc5.1-EGFP-**Valois** | *valois* cDNA | F39-MJ |
| **pAc5.1-mEGFP** (N-ter) EcoRV) (T6-MJ) (Jeske *et al*, 2017) | pAc5.1-mEGFP-**Smo 556-1036** | *Drosophila* ovarian cDNA | JK242 |
| | pAc5.1-mEGFP-**Me31B** | *Drosophila* ovarian *cDNA* | JM145 |
| | pAc5.1-mEGFP-**Trailer Hitch (Tral)** | *Drosophila* ovarian *cDNA* | JM162 |
| | pAc5.1-mEGFP-**Belle** | *belle* cDNA | G2-MJ |
| | pAc5.1-mEGFP-**Ago1** | cDNA | JM171 |
| | pAc5.1-mEGFP-**Ago2** | *Drosophila* ovarian *cDNA* | JM161 |
| | pAc5.1-mEGFP-**Ago3** | *Drosophila* ovarian *cDNA* | JM60 |
| **pAc5.1-mEGFP** (C-ter) (FspAI) (EB02) | pAc5.1-**Smoothened**-mEGFP | *Drosophila* ovarian cDNA | JK192 |
| | pAc5.1-**Smoothened Δ970-1003** -mEGFP | pAc5.1-Smoothened-mEGFP (JK192) | JK214 |
| **pAc5.1** (xxx) (T4-MJ) | pAc5.1-**Long Oskar**-mCherry | *oskar* cDNA | H1-MJ |
| **pAc5.1-mCherry** (EcoRV) (T7-MJ) (Jeske *et al*, 2017) | pAc5.1-mCherry-**Smaug** | *smaug* cDNA | H28-MJ |
| | pAc5.1-mCherry-**Smaug MUT (S250E/L253E)** | Deletion of PH domain from pAc5.1-PH-Smaug MUT (S250E/L253E) (JM265) | JM268 |
| | pAc5.1-mCherry-**Smaug Δ1-70** | pAc5.1-mCherry-**Smaug** (H28-MJ) | JM127 |
| | pAc5.1-mCherry-**Smaug 1-288 Δ156-196** | pAc5.1-mCherry-Smaug 1-288 (H29-MJ) | JK185 |
| | pAc5.1-mCherry-**Smaug Δ1-288** | pAc5.1-mCherry-**Smaug** (H28-MJ) | JM110 |
| | pAc5.1-mCherry-**mouse SAMD4A** | pAc 5.1-PH-mCherry-MmSAMD4A (JK177) | JK183 |
| | pAc5.1-mCherry-**mouse SAMD4B** | pAc 5.1-PH-mCherry-MmSAMD4B (JK181) | JK201 |

| | | | |
|---|---|---|---|
| | pAc5.1-mCherry-**Fused** | *fused* cDNA (LD03657 clone, DGRC) | JK212 |
| | pAc5.1-mCherry-**Fused 1-281** | pAc5.1-PH-mCherry-Fused (JK249) | JK265 |
| | pAc5.1-mCherry-**Fused 282-494** | pAc5.1-PH-mCherry-Fused (JK249) | JK266 |
| | pAc5.1-mCherry-**Fused 495-805** | pAc5.1-PH-mCherry-Fused (JK249) | JK269 |
| **pAc5.1-λN-HA** (EcoRV) (T8-MJ) (gift from Elisa Izaurralde) (Behm-Ansmant *et al*, 2006) | pAc5.1-λN-HA-**eIF4E** | *Drosophila* ovarian cDNA | JM163 |
| **pAc5.1-PH-mEGFP** (FspAI) (JM50) (Salgania *et al*, 2022) | pAc5.1-PH-mEGFP-**Smo 958-1003** | pAc 5.1-mEGFP-Dm Smo 556-1035 (JK168) | JK179 |
| | pAc5.1-PH-mEGFP-**Smo 970-1003** | pAc5.1-PH-mEGFP-Dm Smo 556-1036 (JK178) | JK199 |
| | pAc5.1-PH-mEGFP-**Smo 970-1003 MUT (L978E/L984E/L985E)** | synthetic DNA sequence (IDT) | JK235 |
| | pAc5.1-PH-mEGFP-**Cup** | (Salgania *et al*, 2022) | JM51 |
| | pAc5.1-PH-mEGFP-**mouse Smo 542-793** | *Drosophila* ovarian cDNA | JK193 |
| | pAc5.1-PH-mEGFP-**Smo 556-1036** | pAc 5.1-mEGFP-Dm Smo 556-1035 (JK168) | JK178 |
| | pAc5.1-PH-mEGFP-**Slmb** | *Drosophila* ovarian cDNA | JM251 |
| | pAc5.1-PH-mEGFP-**Bard** | *Drosophila* embryonal cDNA | JK270 |
| | pAc5.1-PH-mEGFP-**14-3-3ζ** | *Drosophila* head cDNA | JK195 |
| | pAc5.1-PH-mEGFP-**14-3-3ε** | *Drosophila* head cDNA | JK196 |
| **pAc5.1-PH-mCherry** (FspAI) (HK49) (Salgania *et al*, 2022) | pAc5.1-PH-mCherry-**Smaug** | *smaug* cDNA | JZ1 |
| | pAc5.1-PH-mCherry-**Smaug 1-281** | *smaug* cDNA | JK243 |
| | pAc5.1-PH-mCherry-**Smaug 1-281 MUT (S250E/L253E)** | Site directed mutagenesis on pAc5.1-PH-mCherry-Smaug 1-281 (JK243) | JM265 |
| | pAc5.1-PH-mCherry-**Smaug 282-595** | *smaug* cDNA | JK244 |
| | pAc5.1-PH-mCherry-**Smaug 596-764** | *smaug* cDNA | JK245 |
| | pAc5.1-PH-mCherry-**Smaug 765-999** | *smaug* cDNA | JK246 |

| | | | |
|---|---|---|---|
| **pAc5.1-OST4-mCherry** (EcoRV) (XH26) (Salgania *et al*, 2022) | pAc5.1-OST4-mCherry-**Short Oskar** (139-606) | (Salgania *et al*, 2022) | H3-MJ |
| **pAc5.1-OST4-EGFP** (FspAI) (JK268) | pAc5.1-OST4-EGFP-**Short Oskar** (139-606) | *oskar* cDNA | JM266 |
| | pAc5.1-OST4-EGFP-**Short Oskar Δ295-352** | Site-directed mutagenesis of pAc5.1-OST4-EGFP-Short Oskar (139-606) (JM266) | JM272 |
| **pMJ-His** (ScaI) (T43-MJ) | pMJ-His-**human SAMD4A 2-156** | synthetic DNA sequence codon-optimized for *E. coli* (IDT) | FH27 |
| | pMJ-His-**human SAMD4A 2-156 H86P** | pMJ-His-human SAMD4A 2-156 (FH27) | JK170 |
| | pMJ-His-**mouse SAMD4A 2-156** | cDNA from mouse P19 cell line (gift from Julien Béthune) | GU45 |
| | pMJ-His-**mouse SAMD4A 2-156 H86P** | pMJ-His-mouse SAMD4A 2-156 (GU45) | GU46 |
| | pMJ-His-**Smaug 37-281** | *smaug* cDNA | JK126 |
| | pMJ-His-**Smaug 70-281** | *smaug* cDNA | JK127 |
| | pMJ-His-**Smaug 70-281 MUT (S250E/L253E)** | Site-directed mutagenesis of pMJ-His-Smaug 70-281 (JK127) | JK257 |
| | pMJ-His-**Smo 970-1003-(GGS)$_4$-Smaug 73-278Δ156-196** | pMJ-His-Smaug 70-278 Δ156-196 (AG4)<br>pAc 5.1-mEGFP-Dm Smo 556-1035 (JK168) | GU44 |
| **pMJ-GST** (SmaI) (T42-MJ) | pMJ-GST-**Smoothened 970-1003** | pAc5.1-PH-mEGFP-Dm Smo 556-1036 (JK178) | JK211 |
| | pMJ-GST-**Smoothened 970-1003 MUT (L978E/L984E/L985E)** | synthetic gene (IDT) | GU48 |
| **pMJ-His-MBP** (ScaI) (T46-MJ) | pMJ-His-MBP-**Oskar 292-325** | pPR3-N-Oskar (N1-MJ) | AG9 |
| **pAc5.1C-Fluc** (EcoRI, XhoI) (E5-MJ) (gift from Elisa Izaurralde) | pAc5.1C-Fluc-**nos 3'UTR** | Luc nos RNA plasmid (Jeske *et al*, 2006) | JK166 |
